# Accurate Nonlinear Mapping between MNI Volumetric and FreeSurfer Surface Coordinate Systems

**DOI:** 10.1101/302794

**Authors:** Jianxiao Wu, Gia H. Ngo, Douglas Greve, Jingwei Li, Tong He, Bruce Fischl, Simon B. Eickhoff, B.T. Thomas Yeo

## Abstract

The results of most neuroimaging studies are reported in volumetric (e.g., MNI152) or surface (e.g., fsaverage) coordinate systems. Accurate mappings between volumetric and surface coordinate systems can facilitate many applications, such as projecting fMRI group analyses from MNI152/Colin27 to fsaverage for visualization, or projecting resting-state fMRI parcellations from fsaverage to MNI152/Colin27 for volumetric analysis of new data. However, there has been surprisingly little research on this topic. Here, we evaluated three approaches for mapping data between MNI152/Colin27 and fsaverage coordinate systems by simulating the above applications: projection of group-average data from MNI152/Colin27 to fsaverage and projection of fsaverage parcellations to MNI152/Colin27. Two of the approaches are currently widely used. A third approach (registration fusion) was previously proposed, but not widely adopted. Two implementations of the registration fusion (RF) approach were considered, with one implementation utilizing the Advanced Normalization Tools (ANTs). We found that RF-ANTs performed the best for mapping between fsaverage and MNI152/Colin27, even for new subjects registered to MNI152/Colin27 using a different software tool (FSL FNIRT). This suggests that RF-ANTs would be useful even for researchers not using ANTs. Finally, it is worth emphasizing that the most optimal approach for mapping data to a coordinate system (e.g., fsaverage) is to register individual subjects directly to the coordinate system, rather than via another coordinate system. Only in scenarios where the optimal approach is not possible (e.g., mapping previously published results from MNI152 to fsaverage), should the approaches evaluated in this manuscript be considered. In these scenarios, we recommend RF-ANTs (https://github.com/ThomasYeoLab/CBIG/tree/master/stable_projects/registration/Wu2017_RegistrationFusion).

## Introduction

Most neuroimaging studies register their participants to a common coordinate system for group analyses (Talairach et al., 1967; Talairach and Tournoux, 1988; Evans et al., 1993; Thompson et al., 1997; Fischl et al., 1999b; Van Essen, 2002). Even studies focusing on individual-specific analyses map individual participants to a common coordinate system (e.g., Gordon et al., 2017), allowing for comparisons across participants or studies. There are two main types of coordinate systems: volumetric and surface. The advantage of volumetric coordinate systems is that both cortical and subcortical structures are represented, in contrast to surface coordinate systems that only focus on the cerebral cortex. Conversely, surface-based coordinate systems allow for more accurate inter-subject registration by respecting the 2D topology of the cerebral cortex (Fischl et al., 1999a; Goebel et al., 2006; Anticevic et al., 2008; Cointepas et al., 2010; Ghosh et al., 2010; Pantazis et al., 2010; Van Essen et al., 2012; Tucholka et al., 2012).

The most popular volumetric coordinate system is the MNI152 template, obtained by group-wise registration of 152 participants (Mazziotta et al., 1995, 2001; Good et al., 2001; Fonov et al., 2011; Grabner et al., 2006). Another common volumetric coordinate system is the single-subject MNI template (i.e., Colin27; Holmes et al., 1998), often used in the neuroimaging software packages SPM and MRIcron for lesion-symptom mapping (Ashburner and Friston, 1999; Rorden et al., 2007). The most popular surface coordinate system is FreeSurfer fsaverage template (Fischl et al., 1999b; Bar and Aminoff, 2003; Filimon et al., 2007; Yeo et al,. 2010a). An important issue with multiple coordinate systems is that results reported in one coordinate system cannot be easily translated to another coordinate system.

While there have been tremendous research efforts on mapping data from individual subjects into common coordinate systems (Collins et al., 1994; Woods et al., 1998; Rueckert et al., 1999; Hellier et al., 2003; Andersson et al., 2007; Ashburner, 2007; Hamm et al, 2010; Yeo et al., 2010b; Yushkevich et al., 2012; Robinson et al., 2014; Tong et al., 2017; Nenning et al., 2017), there is significantly less work on mappings between coordinate systems (Lancaster et al., 2007; Laird et al., 2010). Accurate mapping between volumetric (e.g. MNI152) and surface (e.g. fsaverage) coordinate systems would be useful for many applications. For example, it is a common practice for researchers to perform group analysis in MNI152 space, and then project the results to fsaverage space for visualization (Liu et al., 2009; Sepulcre et al., 2010; Yeo et al., 2015). As another example, resting-state parcellations estimated in fsaverage or fs_LR surface coordinate systems (Yeo et al., 2011; Gordon et al., 2016; Glasser et al., 2016; Schaefer et al., in press) can be projected to the MNI152 coordinate system for analyzing fMRI data of new subjects registered to the MNI152 template. Finally, a more accurate MNI152-fsaverage mapping would facilitate the comparison of thousands of neuroimaging studies reported in either MNI152 or fsaverage coordinate system.

In this work, we evaluate three approaches (including two implementations of one of the approaches) for mapping between volumetric (MNI152 or Colin27) and surface (fsaverage) coordinate systems. The evaluation utilized simulations mimicking the previously described applications: projection of group-average data from MNI152/Colin27 to fsaverage and projection of surface-based parcellations from fsaverage to MNI152/Colin27. We note that the evaluations are not comparisons of volumetric and surface registrations. Instead, the evaluations served to provide error bounds on different mappings between MNI152/Colin27 and fsaverage coordinate systems and to guide the adoption of best practices.

It is also worth emphasizing that a perfect mapping between volumetric and surface coordinate systems is impossible because of registration errors that become irreversible after group averaging. Therefore, the best way of mapping data to fsaverage is by registering subjects directly to fsaverage (e.g., via the official FreeSurfer recon-all pipeline). Similarly, the best way of mapping data to MNI152/Colin27 is by registering subjects directly to the corresponding volumetric template. The approaches evaluated in this paper should only be considered when the best approach is not possible, e.g., mapping previously published results from MNI152 to fsaverage. Whenever the original data from a subject’s native space are available, one should perform registration between the subject’s native space and the desired coordinate system (fsaverage, MNI152 or Colin27) directly, rather than utilize the approaches evaluated in this paper.

## Methods

### Volumetric and surface templates

The MNI152 coordinate system is created by averaging the MRI scans of 152 participants and affords a higher resolution over the original MNI305 average brain. Here we consider the 1mm asymmetric MNI152 template distributed by the FMRIB Software Library (FSL) version 5.0.8. The template was obtained by the linear and nonlinear registration of 152 T1-weighted images (Grabner et al., 2006).

Although MNI152 is the most commonly used volumetric coordinate system, the inter-subject averaging results in the loss of fine anatomical details. Therefore, some research communities (e.g., neuropsychology) prefer single-subject templates. A commonly used single-subject template is Colin27 (also called the MNI single subject template), which is an average image across 27 scans of one subject (Holmes et al., 1998). We used the 1mm Colin27 template from the Statistical Parametric Mappings (SPM) Anatomy Toolbox version 2.2c (Eickhoff et al., 2005).

Finally, the most common surface coordinate system is FreeSurfer fsaverage, which is obtained by spherical alignment of 40 participants (Fischl et al., 1999a, 1999b). As a surface template, fsaverage offers excellent representation of the cortical surface’s intrinsic topological structure as well as multi-scale summary statistics of cortical geometry. It also has an inflated form, which facilitates data visualization. We used the fsaverage template from FreeSurfer version 4.5.0.

### Data and FreeSurfer processing

Data from 1490 subjects from the Brain Genomics Superstruct Project (GSP) were considered (Holmes et al., 2015). All imaging data were collected on matched 3T Tim Trio scanners using the vendor-supplied 12-channel phase-array head coil. Subjects were clinically normal, English-speaking young adults (ages 18 to 35). The structural MRI data consisted of one 1.2mm × 1.2mm × 1.2mm scan for each participant. Details of data collection can be found elsewhere (Yeo et al., 2011; Holmes et al., 2015). The subjects were split into training and test set, each containing 745 subjects.

A second dataset consisted of 30 healthy young adults from the Hangzhou Normal University of the Consortium for Reliability and Reproducibility (CoRR-HNU) dataset (Zuo et al., 2014; Chen et al., 2015). All anatomical images were collected on matched 3T GE Discovery MR750 scanners using an 8-channel head coil. Ten 1.0mm × 1.0mm × 1.0mm scans were performed for each subject across one month. In this paper, we utilized all 10 sessions for all 30 subjects, giving rise to a total of 300 sessions.

The T1 images of the GSP dataset has been previously processed (Holmes et al., 2015) using FreeSurfer 4.5.0 recon-all procedure (http://surfer.nmr.mgh.harvard.edu; Dale et al., 1999; Ségonne et al., 2004, 2007; Fischl et al., 1999a, 1999b, 2001). For consistency, the T1 images of the CoRR-HNU dataset were also processed using the same FreeSurfer version. FreeSurfer constitutes a suite of automatic algorithms that extract models of most macroscopic human brain structures from T1 MRI data. There are three outputs of the recon-all procedure that were important for subsequent analyses.

First, FreeSurfer automatically reconstructs surface mesh representations of the cortex from individual subjects’ T1 images. The cortical surface mesh is inflated into a sphere, and registered to a common spherical coordinate system that aligned the cortical folding patterns across subjects (Fischl et al., 1999a, 1999b). The outcome of this procedure is a nonlinear mapping between the subject’s native T1 space and fsaverage surface space.

Second, the recon-all procedure generates corresponding volumetric (aparc.a2009s+aseg.mgz) and surface (lh.aparc.a2009s.annot and rh.aparc.a2009s.annot) parcellations of 74 sulci and gyri for each subject (Fischl et al., 2004b; Desikan et al., 2006; Destrieux et al., 2010). FreeSurfer assign these labels based on probabilistic information estimated from a manually labeled training set (Destrieux atlas), as well as geometric information derived from the cortical model of the subject. These anatomical segmentations will be utilized in our evaluation of various algorithms for mapping between MNI152/Colin27 and fsaverage.

Third, the recon-all procedure performs a joint registration-segmentation procedure that aligns the T1 image to an internal FreeSurfer volumetric space^1^, while classifying each native brain voxel into one of multiple brain structures, such as the thalamus and caudate (Fischl et al., 2004a, 2004b). The outcome of this procedure is a nonlinear mapping between the subject’s native T1 space and FreeSurfer internal volumetric space. The nonlinear mapping is represented by a dense displacement field (i.e., a single displacement vector at each 2-mm isotropic atlas voxel) and can be found in the file “talairach.m3z” (under the “mri/transforms” folder of the recon-all output).

### Affine and MNIsurf

Two existing approaches (Affine and MNIsurf) for mapping between MNI152 and fsaverage coordinate systems were identified. Both approaches have been discussed on the FreeSurfer mailing list and might be considered as “recommended” FreeSurfer approaches.

Figure 1 summarizes the Affine approach for mapping between MNI152 and fsaverage surface coordinate systems. The Affine approach made use of an affine transformation between the MNI152 template and fsaverage volume space (Figure 1A) provided by FreeSurfer (i.e., $FREESURFER_HOME/average/mni152.register.dat). This affine transformation can be concatenated with the mapping between fsaverage volume and fsaverage surface (Figure 1B) using FreeSurfer functions (mri_vol2surf and mri_surf2vol), thus yielding a mapping between MNI152 and fsaverage coordinate systems.

**Figure 1.**
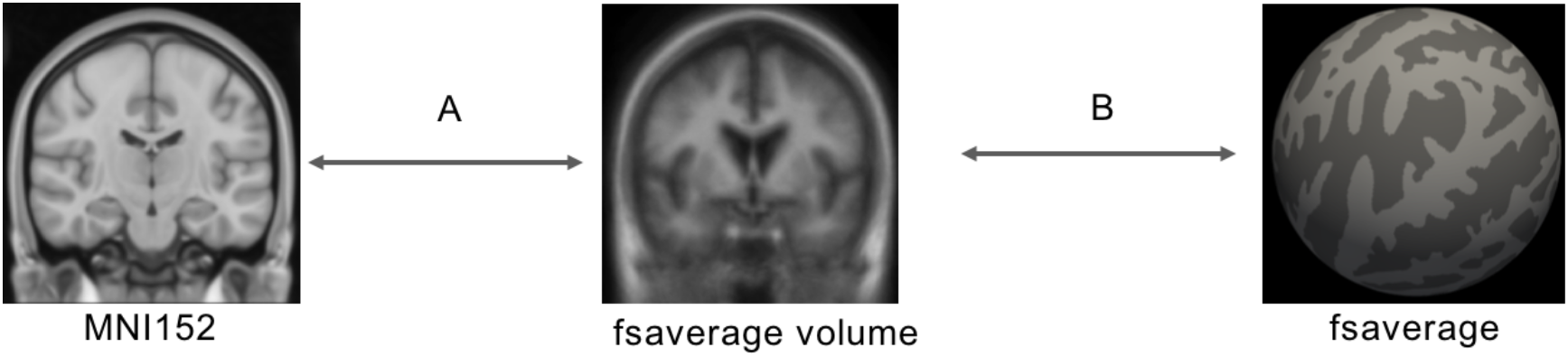
Affine procedure. (A) MNI152 and fsaverage volume was aligned using an affine transformation. (B) FreeSurfer provides a mapping between fsaverage volume and fsaverage surface. Concatenating the two transformations result in a mapping between MNI152 and fsaverage surface.

One drawback of this approach is that an affine transformation is unlikely to eliminate nonlinear anatomical differences between MNI152 and fsaverage volume. Simply replacing the affine transformation with a nonlinear warp (Van Essen et al., 2012) might not be helpful because the fsaverage volume is a blurry average of 40 subjects after affine registration; fine anatomical details have already been lost.

Figure 2 summarizes the MNIsurf approach for mapping between MNI152 and fsaverage surface coordinate systems. The MNI152 template was first processed with FreeSurfer recon-all. The recon-all process involved extracting MNI152 template’s cortical ribbon and reconstructing the cortical surface (Figure 2A). FreeSurfer commands (mri_vol2surf and mri_surf2vol) could then be utilized to map between MNI152’s cortical ribbon (as segmented by recon-all) and fsaverage surface (Figure 2B).

**Figure 2.**
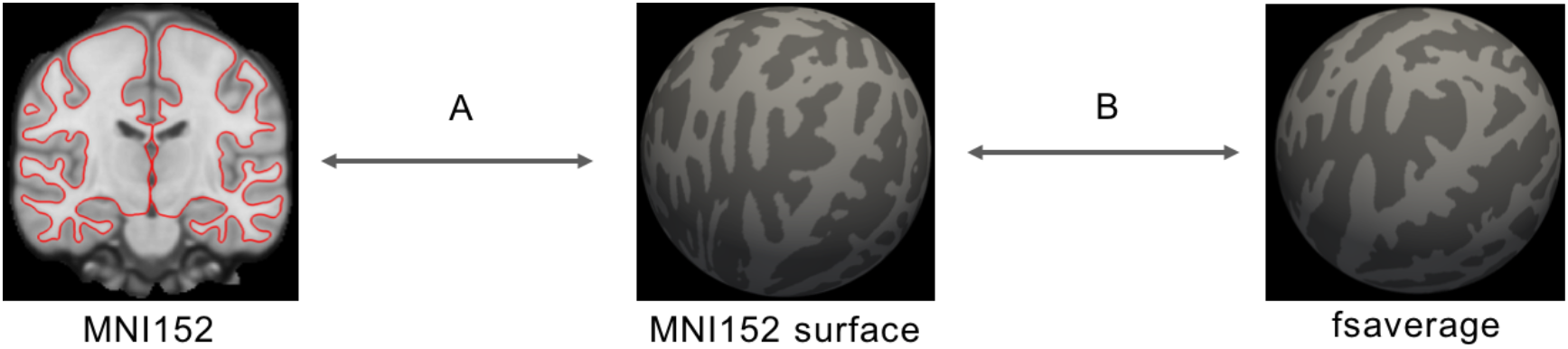
MNIsurf procedure. The MNI152 template was processed using FreeSurfer recon-all. The cortical ribbon of MNI152 was (A) extracted and (B) aligned to fsaverage surface during the recon-all procedure.

One drawback of MNIsurf is that the cortical ribbon of a typical subject mapped to MNI152 coordinate system will not exactly match the group-average MNI152 cortical ribbon (which is abnormally thin and misses some low-frequency and/or thin folds due to inter-subject averaging). Consequently, there will be irreversible registration errors from averaging subjects mapped to the MNI152 coordinate system. MNIsurf does not take into account these irreversible registration errors because it simply maps the cortical ribbon of MNI152 directly to fsaverage surface.

### Registration fusion: RF-M3Z and RF-ANTs

The registration fusion (RF) approach was first introduced by Buckner and colleagues (Buckner et al., 2011; Yeo et al., 2011). Figure 3 summarizes the original implementation. Recall that by applying FreeSurfer recon-all procedure to each GSP training subject, we have generated for each subject a nonlinear mapping between the subject’s cortical ribbon and fsaverage surface space (Figure 3C) and a nonlinear mapping between the subject’s T1 volume and FreeSurfer internal volumetric space (Figure 3B). By also processing the MNI152 template with FreeSurfer recon-all, we also obtained a nonlinear mapping between the MNI152 template and FreeSurfer internal volumetric space (Figure 3A). By concatenating the three transformations (Figure 3A, Figure 3B and Figure 3C) for each subject, a mapping between MNI152 and fsaverage coordinate systems for each GSP training subject was obtained. By averaging across all 745 training subjects, a final mapping between MNI152 and fsaverage coordinate systems was obtained. This mapping is referred to as RF-M3Z.

**Figure 3.**
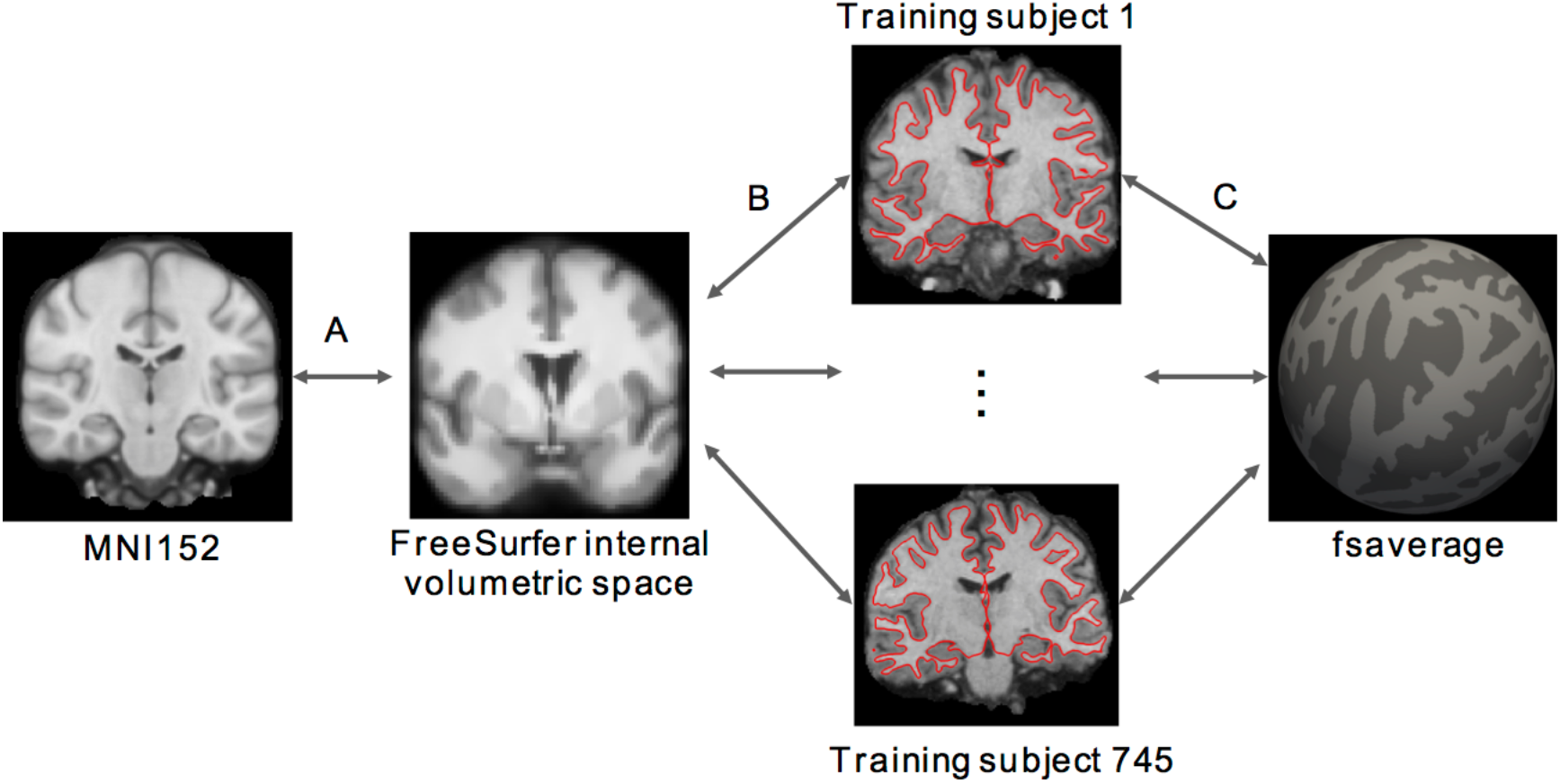
Registration fusion (RF-M3Z) procedure. Each subject’s T1 volume is mapped to the (A, B) MNI152 template and (C) fsaverage surface. By concatenating the mappings for each subject and then averaging the deformations across all 745 training subjects, we created a mapping between MNI152 and fsaverage surface space. All mappings (A, B and C) were generated using FreeSurfer’s recon-all procedure. More specifically, mappings A and B were provided by the talairach.m3z files generated by recon-all, so we refer to the resulting MNI152-fsaverage mapping as RF-M3Z.

Visual inspection suggested that the mappings between MNI152 and individual subjects (concatenations of transformations in Figure 3A and Figure 3B) were of good quality (Buckner et al., 2011; Yeo et al., 2011). However, by concatenating two deformations, small registration errors in each deformation may be compounded to result in large registration errors. Furthermore, FreeSurfer is optimized for processing the brains of individual subjects, not an average brain like the MNI152 template.

Therefore, we also considered a second implementation, where the individual subjects and the MNI152 template were directly registered using ANTs (Avants et al., 2007, 2009). More specifically, each GSP training subject’s T1 image was directly registered to the MNI152 template using an affine transformation followed by Symmetric Normalization (Figure 4A). Like RF-M3Z, the mapping between each subject’s cortical ribbon and fsaverage surface space was provided by FreeSurfer recon-all (Figure 4B). By concatenating the two transformations (Figure 4A and Figure 4B) for each subject, a mapping between MNI152 and fsaverage coordinate systems for each GSP training subject was obtained. By averaging across all 745 training subjects, a final mapping between MNI152 and fsaverage coordinate systems was obtained. This mapping is referred to as RF-ANTs. It is important to note that this does not constitute a comparison of FreeSurfer and ANTs (Klein et al., 2010), since FreeSurfer is not being used in the way it was designed (i.e., individual subject analyses) in the case of RF-M3Z.

**Figure 4.**
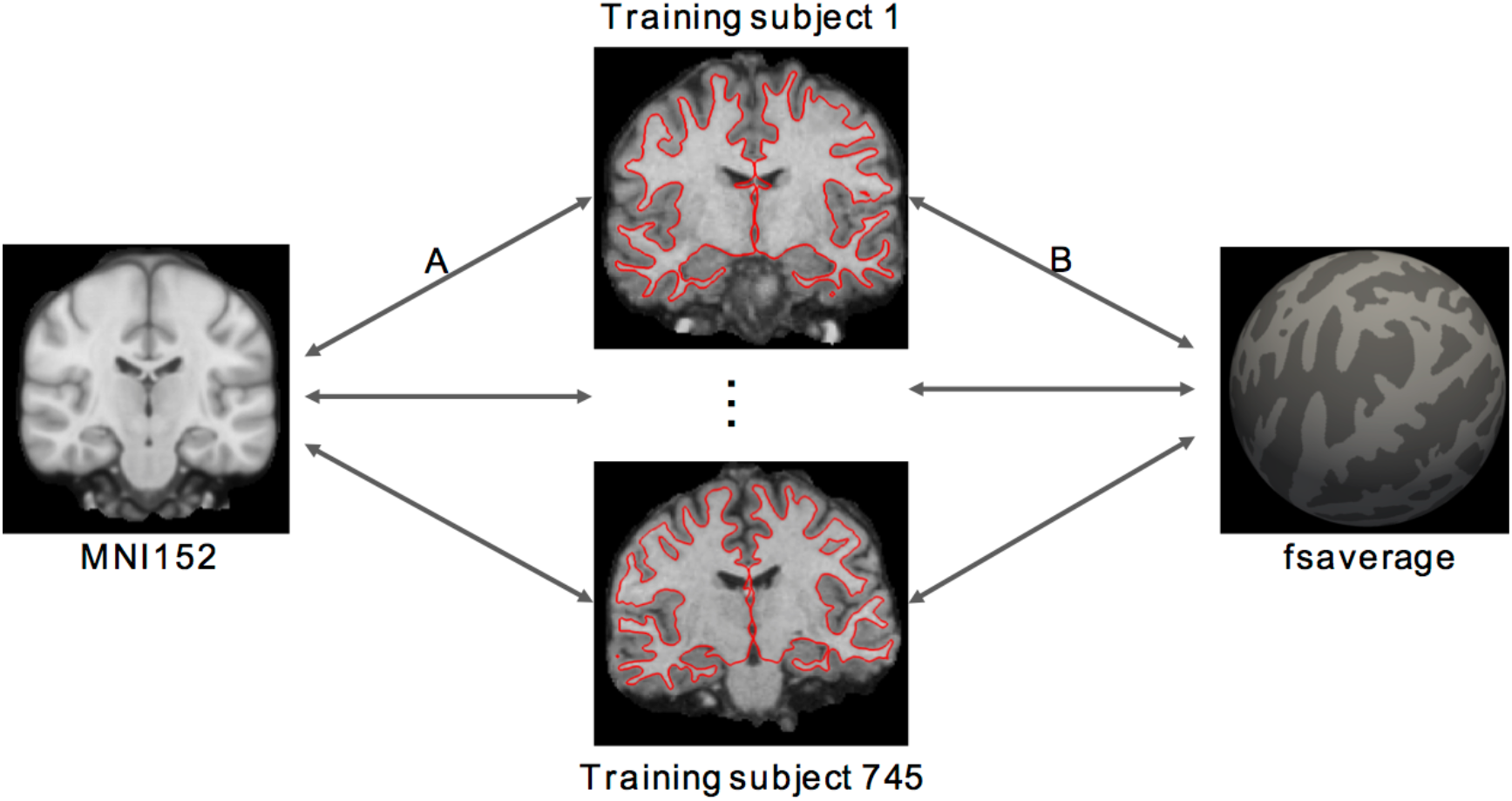
Registration fusion (RF-ANTs) procedure. Each subject’s T1 volume is mapped to the (A) MNI152 template and (B) fsaverage surface. By concatenating the mappings (A and B) for each subject and then averaging the deformations across all 745 training subjects, we created a mapping between MNI152 and fsaverage. Mapping (A) was generated using ANTs, so we refer to the resulting MNI152-fsaverage mapping as RF-ANTs.

### Tight and loose cortical masks for fsaverage-to-MNI152 mappings

It is worth mentioning an important asymmetry in the generation of the MNI-to-fsaverage and fsaverage-to-MNI mappings. When computing the MNI-to-fsaverage mapping, each subject yielded a mapping between *every* fsaverage vertex and *some* MNI location, which allowed for a simple averaging of MNI-to-fsaverage mappings across all 745 training subjects. By contrast, when computing the fsaverage-to-MNI mapping, not every training subject yielded a mapping between *every* MNI location and *some* fsaverage vertex because not every MNI location corresponded to the cerebral cortex of every subject.

Therefore, when computing the fsaverage-to-MNI mapping, we defined two cortical masks. Figure 5 illustrates the two MNI152 masks and the difference between them. The tight cortical mask corresponded to the cortex for at least 50% of the subjects (Figure 5A), while the loose cortical mask corresponded to the cortex for at least 15% of the subjects (Figure 5B). For each tight cortical mask voxel (Figure 5A), the fsaverage-to-MNI152 mappings were averaged across all subjects with valid fsaverage-to-MNI152 mappings for the voxel. The averaged mapping was then grown outwards to fill the entire loose cortical mask. More specifically, for each voxel outside the tight mask (but within the loose mask; Figure 5C), its nearest voxel within the tight mask (Figure 5A) was identified based on Euclidean distance. The voxel was then assigned the same fsaverage surface coordinates as its nearest voxel within the tight mask. Therefore, fsaverage surface data can be projected to fill up the entire loose cortical mask in the MNI152 template. This procedure was repeated for Affine, MNIsurf, RF-M3Z and RF-ANTs.

**Figure 5.**
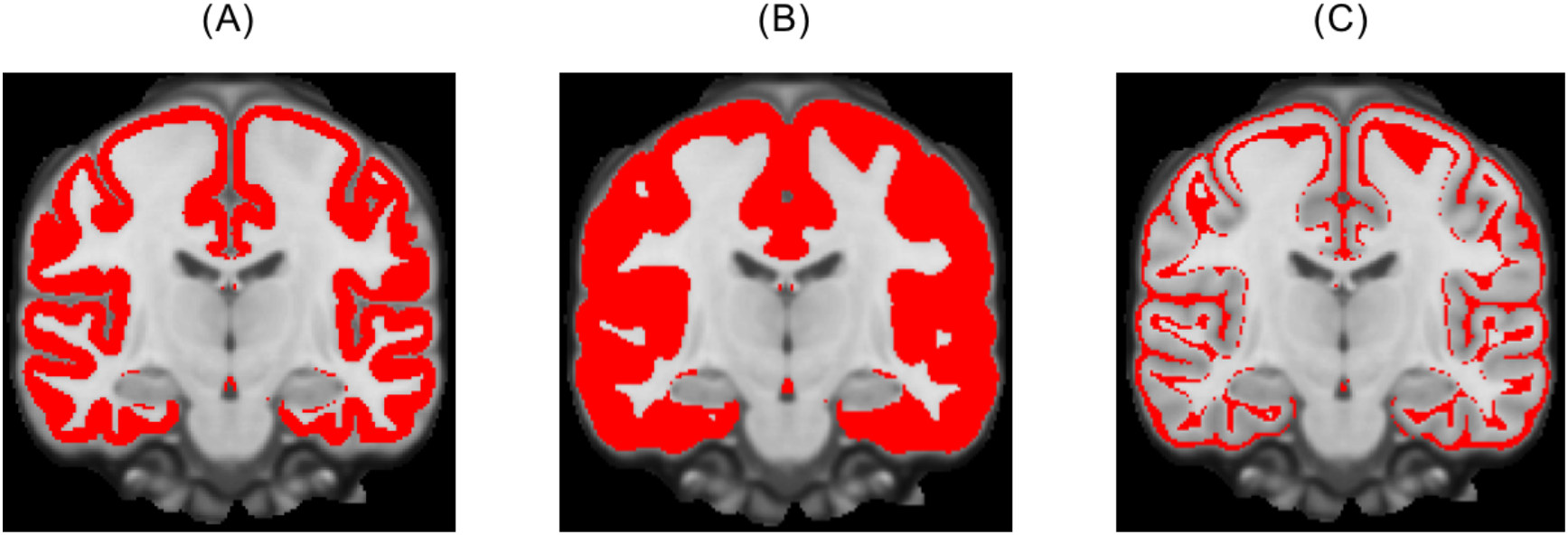
Cortical masks for fsaverage-to-MNI152 mappings. (A) Tight cortical mask corresponding to 50% of the 745 GSP training subjects. (B) Loose cortical mask corresponding to 15% of the training subjects. (C) Difference between tight and loose cortical masks.

### MNI152-to-fsaverage evaluation

To evaluate the MNI152-to-fsaverage projection, let’s consider a possible usage scenario. Researchers often project data (e.g., fMRI) from subjects’ native spaces to MNI152 coordinate system for some form of group analysis. The outcome of the group analysis can be visualized in the volume, but is often projected to fsaverage surface for visualization. By contrast, data from subjects’ native space can be directly projected to fsaverage surface for group analysis. The subjects-to-MNI152-to-fsaverage results should ideally be close to the subjects-to-fsaverage results.

To simulate the above scenario, recall that we have processed the 745 GSP test subjects using FreeSurfer recon-all, yielding corresponding surface (lh.aparc.a2009s.annot and rh.aparc.a2009s.annot) and volumetric (aparc.a2009s+aseg.mgz) parcellations of 74 sulci and gyri per cortical hemisphere (i.e., Destrieux parcellation). Figure 6A illustrates the superior temporal sulcus label in two GSP test subjects. The parcellation labels were projected to MNI152 coordinate system using ANTs and averaged across subjects, resulting in a ANTs-derived volumetric probabilistic map per anatomical structure. The probabilistic maps simulated the group-average results from typical fMRI studies. As an example, Figure 6B illustrates the ANTs-derived MNI152 volumetric probabilistic

**Figure 6.**
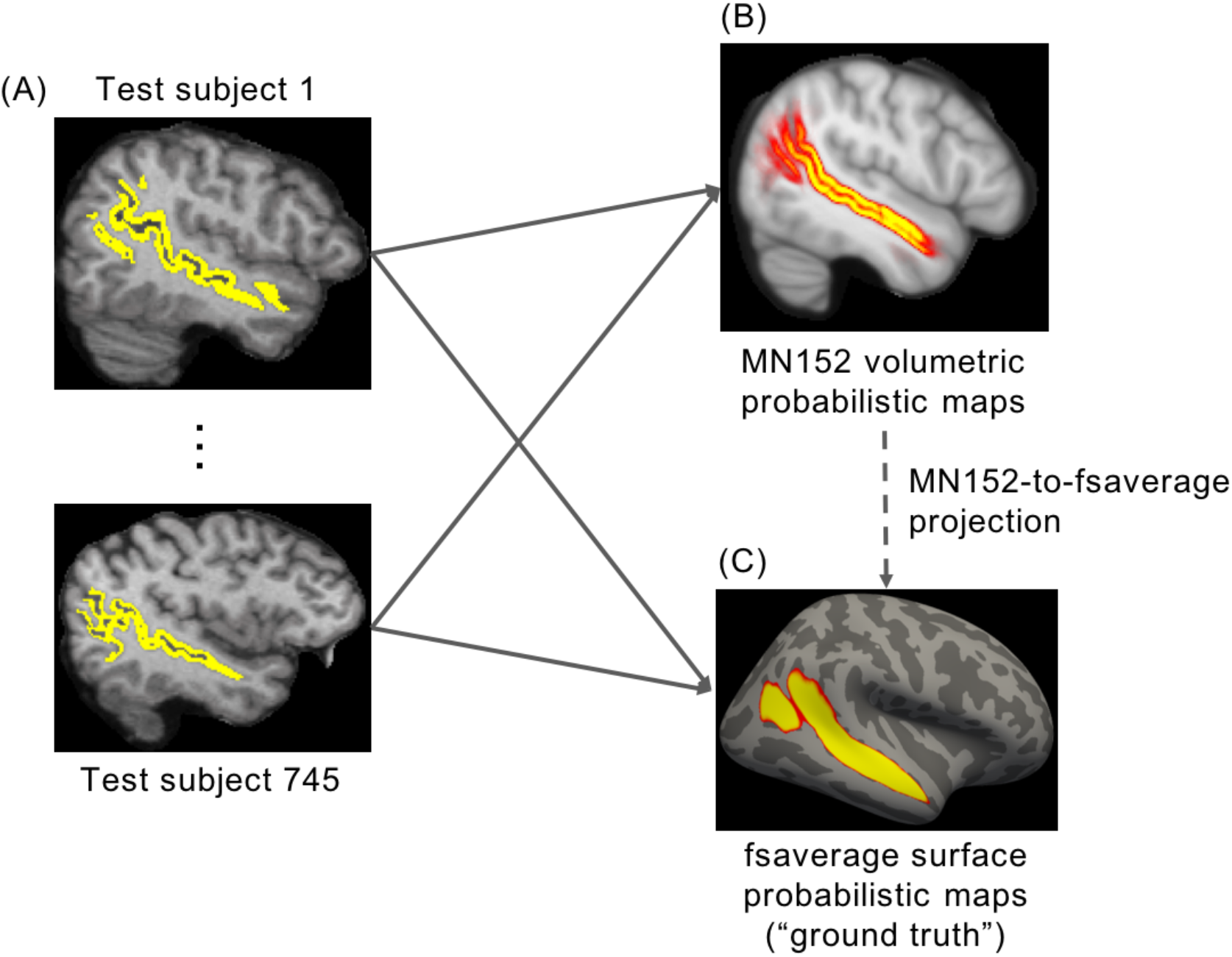
MNI152-to-fsaverage evaluation. (A) Parcellation labels from each subject were projected to (B) MNI152 and (C) fsaverage. The projected labels were averaged across subjects, resulting in a probabilistic map per anatomical structure in (B) MNI152 and (C) fsaverage respectively. Figure shows superior temporal sulcus as an example. The latter maps in (C) fsaverage were used as “ground truth”. The MNI152 probabilistic maps can then be projected to fsaverage surface using the various projection approaches (dotted arrow) for comparison with the “ground truth” maps. map of the superior temporal sulcus.

The MNI152 volumetric probabilistic maps (Figure 6B) can then be projected to fsaverage surface using the various MNI152-to-fsaverage projection approaches (dotted arrow in Figure 6) for comparison with “ground truth” surface probabilistic maps (Figure 6C). The “ground truth” surface probabilistic maps were obtained by averaging the surface parcellations across subjects in fsaverage surface space, mapped from each subject using FreeSurfer. As an example, Figure 6C shows the “ground truth” surface probabilistic map of the superior temporal sulcus.

To quantify the disagreement between the projected probabilistic map and the “ground truth” surface probabilistic map of an anatomical structure, the Normalized Absolute Difference (NAD) metric was used. The NAD metric was defined as the absolute difference between the two maps, summed across all vertices and divided by the sum of the “ground truth” probabilistic map. This metric measured the dissimilarity between the two maps, normalizing for the size of the anatomical structure. A lower NAD value indicates better performance.

For every pair of approaches, the NAD metric for each of the 74 anatomical structures were averaged between the two hemispheres and submitted to a paired-sample t-test. Multiple comparisons were corrected using a false discovery rate (FDR; Benjamini and Hochberg, 1995) of q < 0.05. All the p values reported in subsequent sections survived the false discovery rate.

### fsaverage-to-MNI152 evaluation

To evaluate the fsaverage-to-MNI152 projection, let’s consider a possible usage scenario. It is unlikely that researchers would directly project individual subjects’ fMRI data onto fsaverage surface space for group-level analysis, and then project their results into MNI152 space for visualization. A more likely scenario might be the projection of surface-based resting-state fMRI cortical parcellations (Yeo et al., 2011; Gordon et al., 2015; Glasser et al., 2016; Schaefer et al., in press) to MNI152 space. The projected resting-state fMRI parcellation can then be utilized for analyzing new data from individual subjects registered to the MNI152 coordinate system. In this scenario, it would be ideal if the projected fsaverage-to-MNI152 resting-state parcellation were the same as a parcellation that was estimated from resting-state fMRI data directly registered to MNI152 space.

To simulate the above scenario, the Destrieux anatomical parcellation of each GSP test subject (Figure 7A) was projected to fsaverage and combined into a winner-takes-all parcellation (Figure 7B). The surface-based parcellation can then be projected to MNI152 using the various fsaverage-to-MNI152 projection approaches (dotted arrow in Figure 7) for comparison with the “ground truth” volumetric parcellation (Figure 7C). The “ground truth” volumetric parcellation was obtained by projecting the individual subjects’ anatomical parcellations (Figure 7A) into MNI152 space (using ANTs) and then combined into a winner-take-all parcellation (Figure 7C).

**Figure 7.**
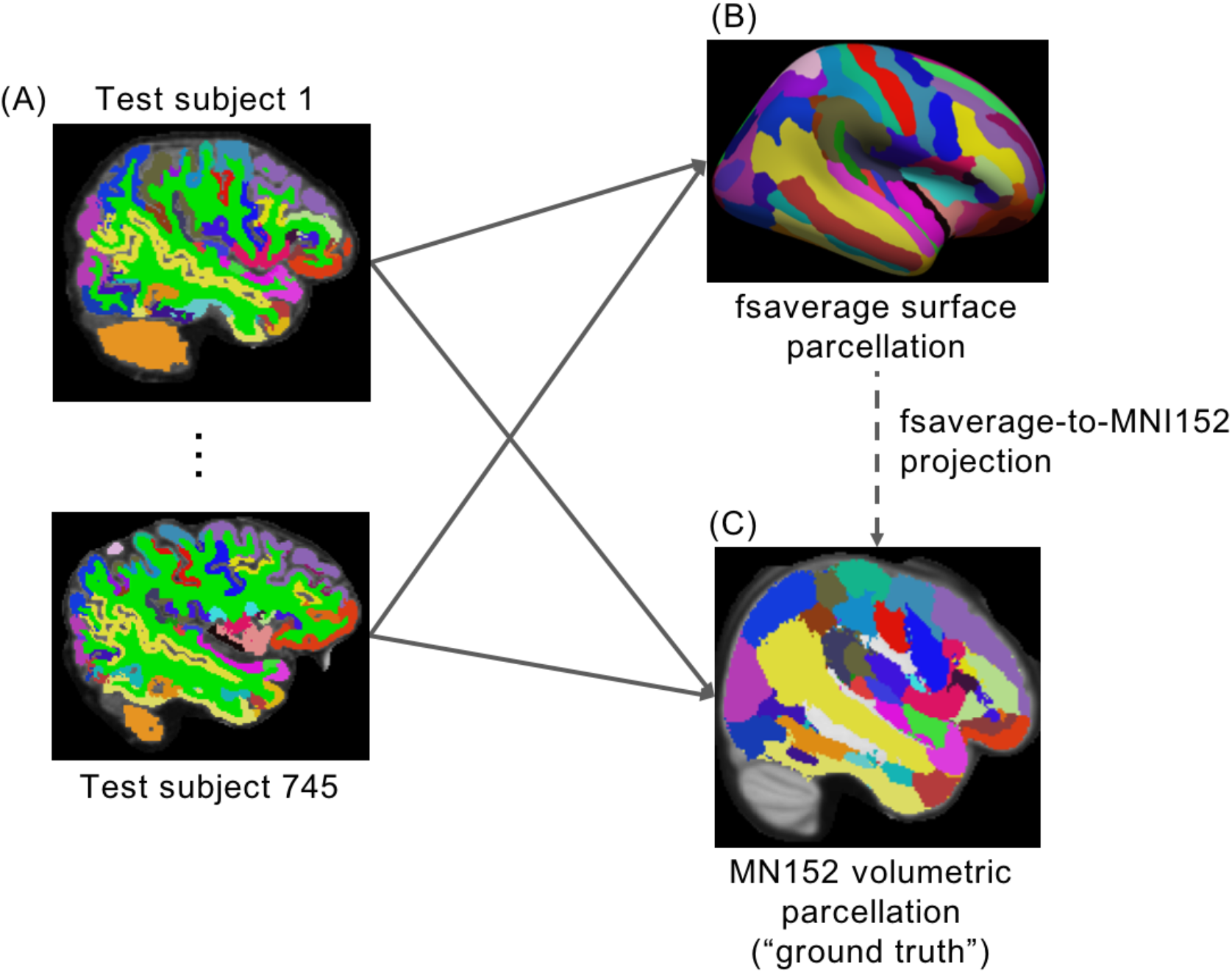
Fsaverage-to-MNI152 evaluation. (A) Parcellation from each subject was projected to (B) fsaverage and (C) MNI152. By combining the parcellations across subjects, winner-takes-all parcellations were obtained in (B) MNI152 and (C) fsaverage respectively. The latter was used as “ground truth”. The fsaverage winner-takes-all parcellation can be projected to MNI152 coordinate system using the various projection approaches (dotted arrow) for comparison with the “ground truth” MNI152 parcellation.

To quantify the agreement between the projected parcellation and the “ground truth” parcellation, the Dice coefficient was computed for each of the 74 anatomical regions per hemisphere. A higher Dice value indicates better performance.

For every pair of approaches, the Dice metric for each of the 74 anatomical structures were averaged between the two hemispheres and submitted to a paired-sample t-test. Multiple comparisons were corrected using a false discovery rate (FDR; Benjamini and Hochberg, 1995) of q < 0.05. All the p values reported in subsequent sections survived the false discovery rate.

### Generalization to new data (CoRR-HNU) and FSL FNIRT

The RF mappings were derived using the GSP training set. To ensure the mappings generalize to new data, the above evaluations (MNI152-to-fsaverage and fsaverage-to-MNI152) were repeated using the CoRR-HNU dataset. Furthermore, the previous evaluation procedures utilized ANTs to project subjects’ anatomical parcellations to MNI152 (Figure 6B and Figure 7C), resulting in possible biases in favor of RF-ANTs. As such, the above evaluations were repeated using FSL FLIRT/FNIRT (Andersson et al., 2007). More specifically, FLIRT/FNIRT was utilized to project individual subjects’ parcellation to MNI152 space to obtain FNIRT-derived CoRR-HNU MNI152 volumetric probabilistic maps (Figure 6B), as well as FNIRT-derived CoRR-HNU MNI152 winner-take-all parcellation (Figure 7C).

### Registration fusion convergence

In the previous analyses, as many training subjects as available (N=745) were used to construct the average mappings for the RF approaches. Here, we investigated the relationship between the accuracy of the RF approaches and the number of subjects used. More specifically, the MNI152-to-fsaverage evaluation (using ANTs-derived GSP MNI152 maps) were repeated using RF mappings averaged across different number of subjects.

### Colin27-to-fsaverage and fsaverage-to-Colin27

The previous mappings and evaluations were repeated for Colin27. In the case of the Affine approach, FreeSurfer does not provide a corresponding Colin27-to-fsaverage-volume warp. Therefore, an affine warp was generated using FSL FLIRT.

Since we are now working with the Colin27 template, the MNIsurf approach was renamed as Colin27surf. It should be noted that unlike that of the MNI152 template, the cortical ribbon of the Colin27 template is not abnormally thin (since it is a single subject template). However, using a single subject prevents the use of cross-subject variance measures that can stabilize inter-subject registration (Fischl, 1999b). Therefore, we also expect registration errors between the cortical ribbon of a typical subject and Colin27. Consequently, Colin27surf does not take into account irreversible registration errors because it simply maps the cortical ribbon of Colin27 directly to fsaverage.

## Results

### MNI152-to-fsaverage projection

Figure 8 shows the projection of ANTs-derived MNI152 probabilistic maps of four representative anatomical structures to fsaverage surface space for the GSP test set. Figure 9 shows the projection of FNIRT-derived MNI152 probabilistic maps of four representative anatomical structures to fsaverage surface space for the CoRR-HNU dataset. The black boundaries correspond to the winner-takes-all parcellation obtained by thresholding the “ground truth” GSP or CoRR-HNU fsaverage surface probabilistic maps respectively. Visual inspection of Figures 8 and 9 suggests that the projected probabilistic maps corresponded well to the “ground truth” for all approaches, although there was also clear bleeding to adjacent anatomical structures for the central sulcus and middle frontal sulcus.

**Figure 8.**
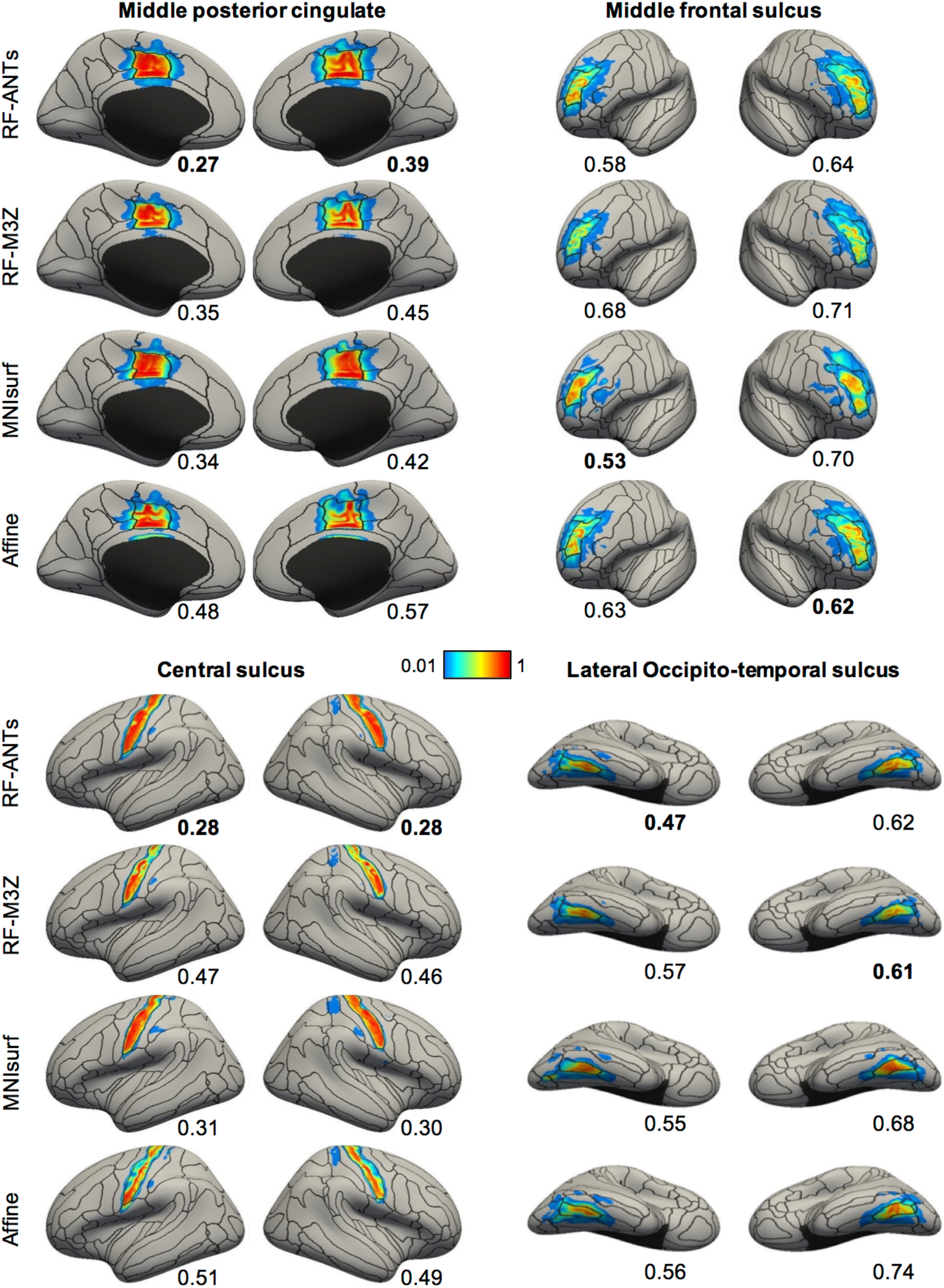
Visualization of ANTs-derived MNI152 probabilistic maps projected to fsaverage surface space in the GSP test set. Four representative structures are shown. Black boundaries correspond to the “ground truth” winner-takes-all parcellation. The value below each cortical surface shows the Normalized Absolute Difference (NAD) between projected probabilistic map and “ground truth” probabilistic map, where a smaller value indicates better performances. Best NAD for each region is **bolded**.

**Figure 9.**
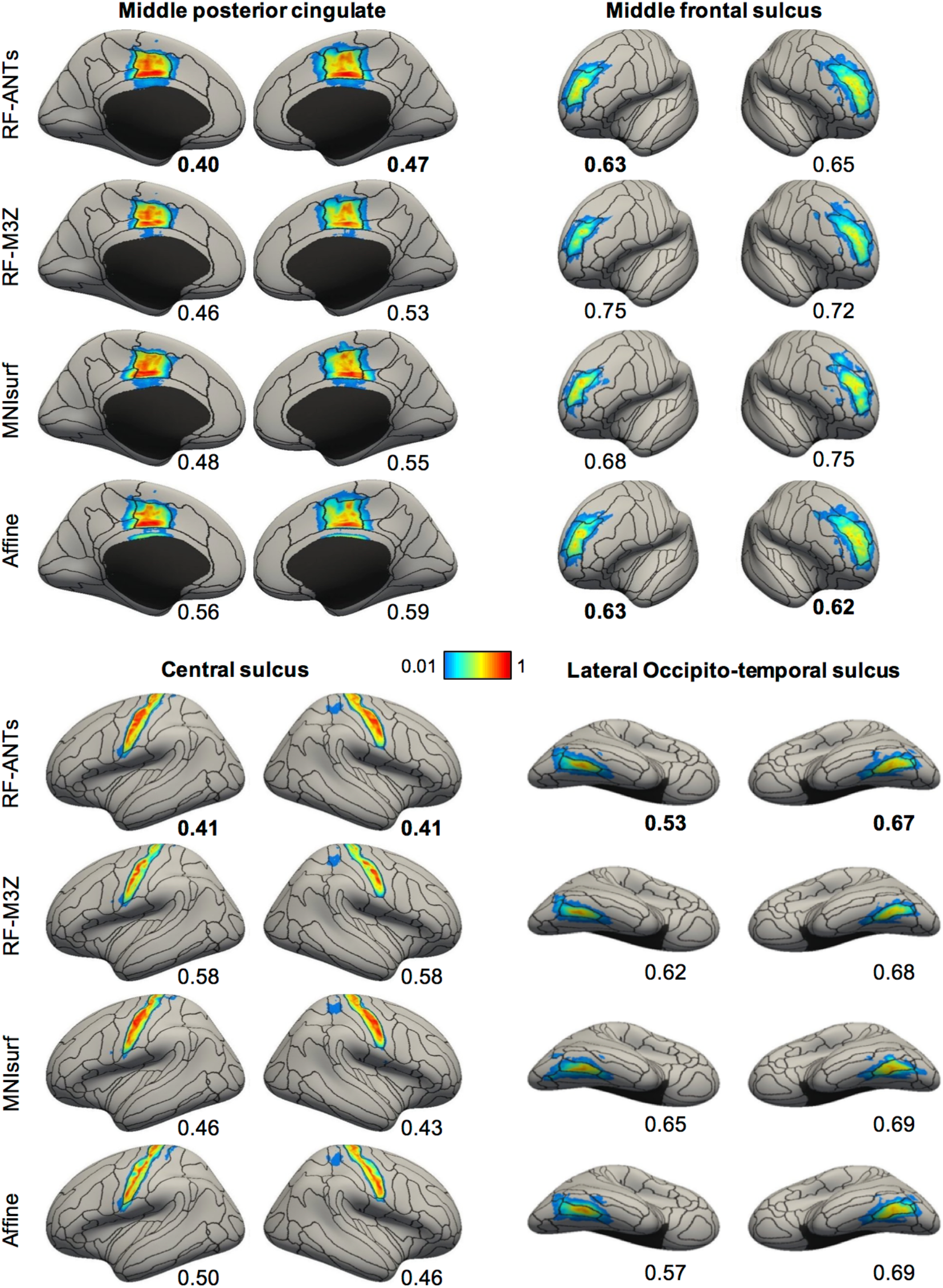
Visualization of FNIRT-derived MNI152 probabilistic maps projected to fsaverage surface space in the CoRR-HNU dataset. Four representative structures are shown. Black boundaries correspond to the “ground truth” winner-takes-all parcellation. The value below each cortical surface shows the Normalized Absolute Difference (NAD) between projected probabilistic map and “ground truth” probabilistic map, where a smaller value indicates better performances. Best NAD for each region is **bolded**.

The NAD evaluation metric is shown below each brain in Figures 8 and 9. A lower value indicates closer correspondence with the “ground truth” probabilistic map. The NAD generally agreed with the visual quality of the projections, suggesting its usefulness as an evaluation metric. For example, in Figure 8, the projection of the left ANTs-derived middle posterior cingulate probabilistic map using RF-ANTs visually matched the “ground truth” black boundaries very well, resulting in a low NAD of 0.27. On the other hand, the corresponding projection using MNIsurf aligned well with the posterior, but not the anterior, portion of the “ground truth” black boundaries, resulting in a worse NAD of 0.34 (Figure 8).

Figure 10 shows the NAD metric averaged across all anatomical structures within each hemisphere. When ANTs-derived GSP MNI152 probabilistic maps were used, RF-ANTs was the best (p < 0.01 corrected). RF-M3Z and MNIsurf showed comparable performance and were both significantly better than Affine (p < 0.01 corrected). When FNIRT-derived CoRR-HNU MNI152 probabilistic maps were used, RF-ANTs was also the best (p < 0.01 corrected). RF-M3Z and Affine showed comparable performance and were both significantly better than MNIsurf (p < 0.04 corrected and p < 0.02 corrected). To summarize, RF-ANTs always performed the best. We note that hemispheric differences within each approach were not statistically significant (all p > 0.2).

**Figure 10.**
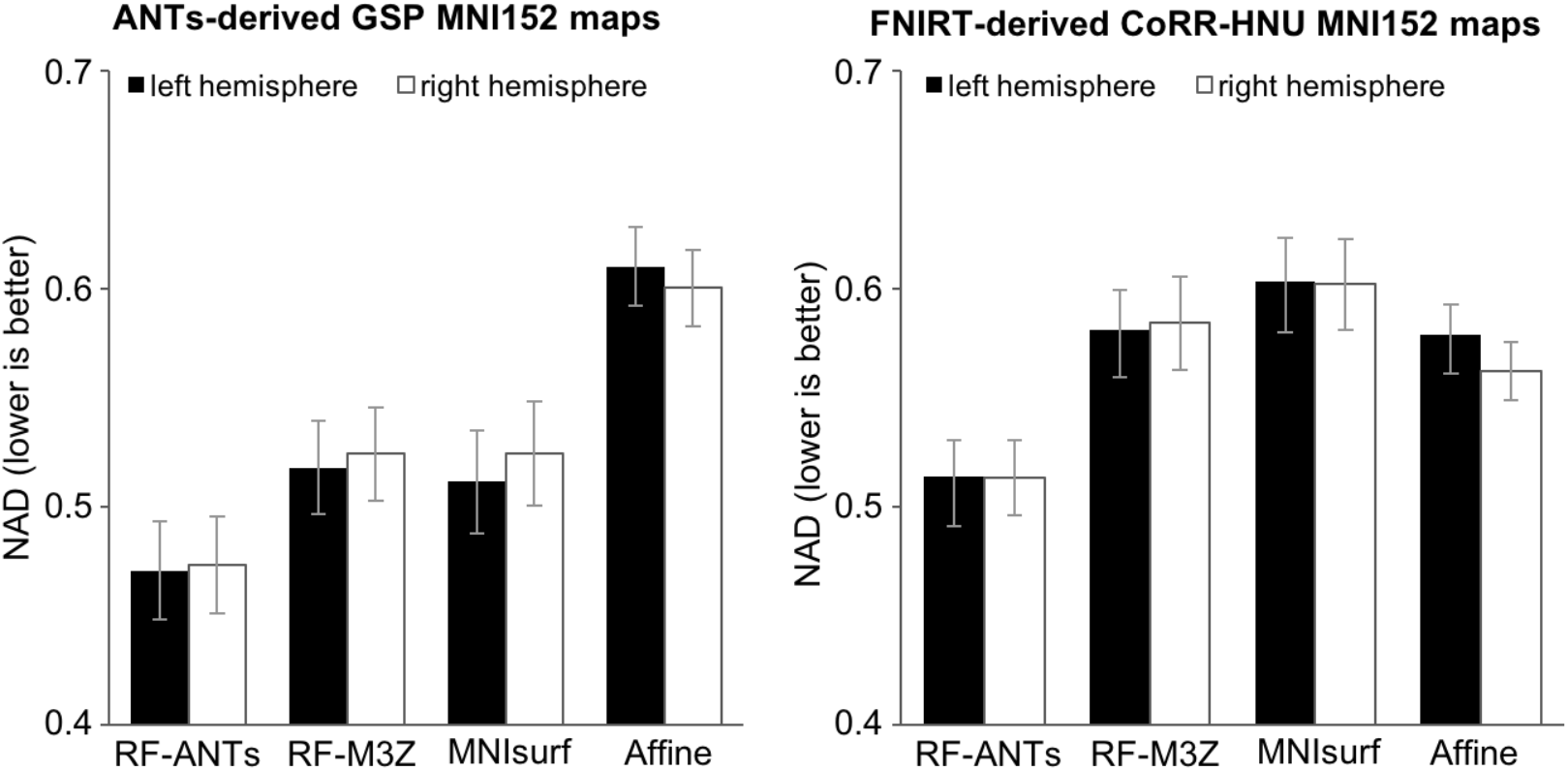
Normalized Absolute Difference (NAD) of MNI152 probabilistic maps projected to fsaverage surface space. (Left) Results for ANTs-derived GSP MNI152 probabilistic maps. (Right) Results for FNIRT-derived CoRR-HNU MNI152 probabilistic maps. The bars represent the NADs averaged across all 74 probabilistic maps within left hemisphere (black) and right hemisphere (white). Error bars correspond to standard errors across the 74 anatomical structures. Overall, RF-ANTs performed the best.

### fsaverage-to-MNI152 projection

Figure 11 illustrates the projection of the fsaverage winner-takes-all parcellation to MNI152 volumetric space for the GSP test set, juxtaposed against black boundaries of ANTs-simulated “ground truth” segmentations. Figure 12 illustrates the projection of the fsaverage winner-takes-all parcellation to MNI152 volumetric space for the CoRR-HNU dataset, juxtaposed against black boundaries of FNIRT-simulated “ground truth” segmentations. Figures 11A and 12A show the fsaverage-to-MNI152 projections before the dilation within the loose cortical mask (see Methods). Figures 11B and 12B show the fsaverage-to-MNI projections after the dilation, with insets illustrating example regions with obvious differences across methods.

**Figure 11.**
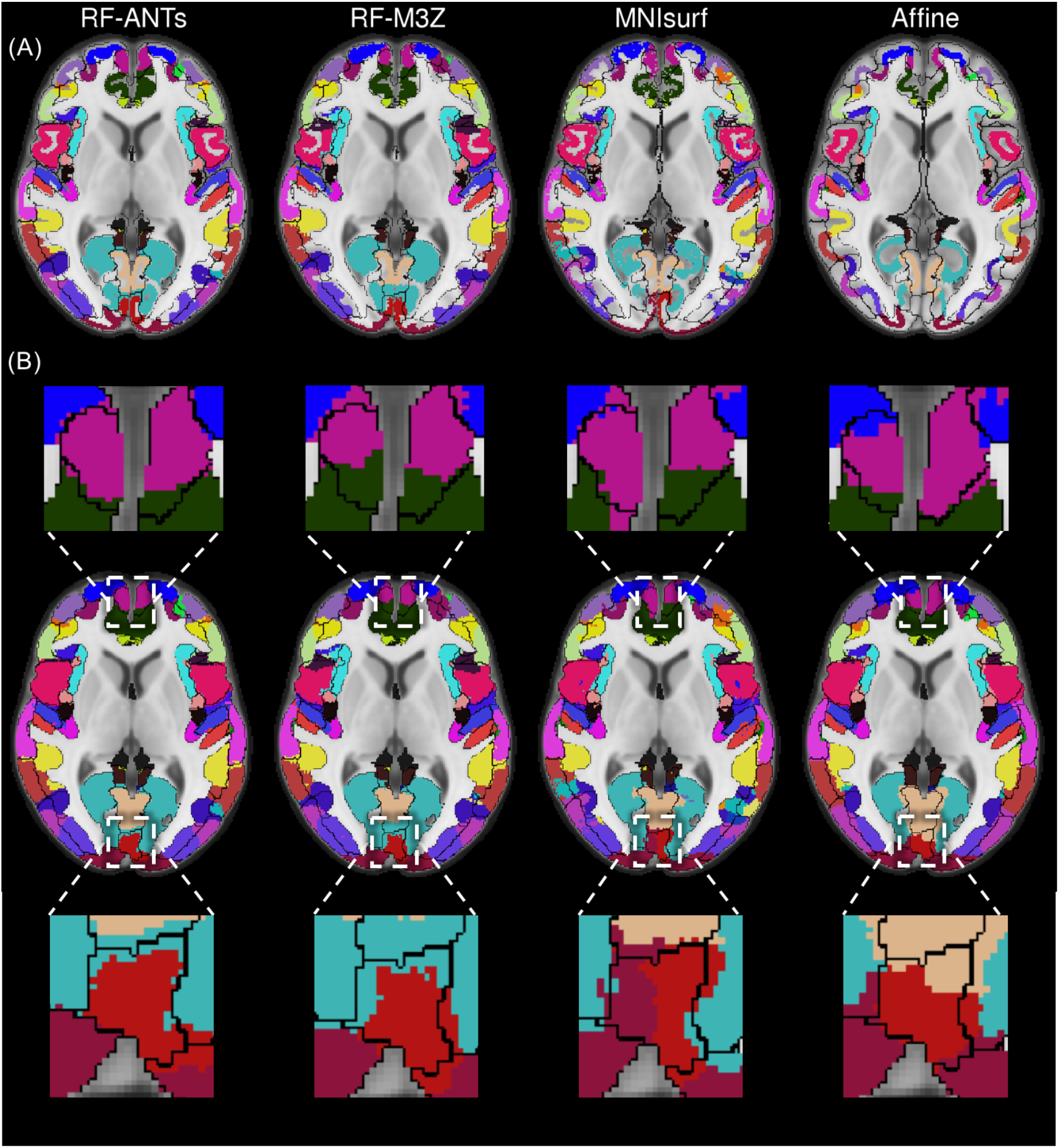
Winner-takes-all fsaverage parcellation projected to MNI152 volumetric space with ANTs-simulated “ground truth” (black boundaries) in the GSP test set. (A) Projections before dilation within loose cortical mask. (B) Projections after dilation within loose cortical mask.

**Figure 12.**
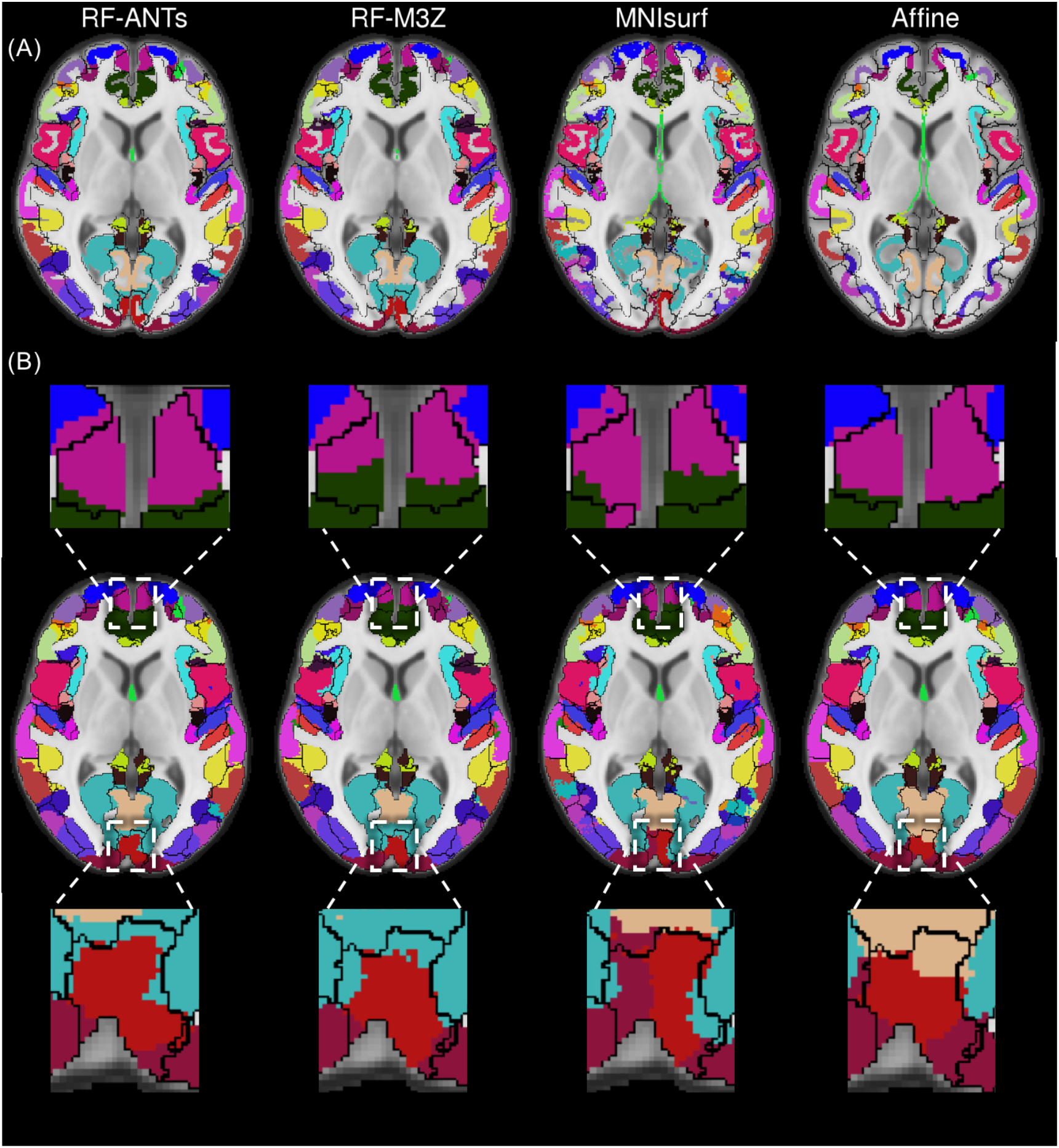
Winner-takes-all fsaverage parcellation projected to MNI152 volumetric space with FNIRT-simulated “ground truth” (black boundaries) in CoRR-HNU dataset. (A) Projections before dilation within loose cortical mask. (B) Projections after dilation within loose cortical mask.

Figure 13 shows the Dice metric averaged across all anatomical structures within each hemisphere. In the case of GSP test set (using ANTs-simulated “ground truth”), RF-ANTs was the best (p < 0.01 corrected). RF-M3Z, MNIsurf and Affine all showed comparable performance. In the case of the CoRR-HNU dataset (using FNIRT-simulated “ground truth”), RF-ANTs and Affine were the best (p < 0.01 corrected). There was no statistically significant difference between RF-ANTs and Affine.

**Figure 13.**
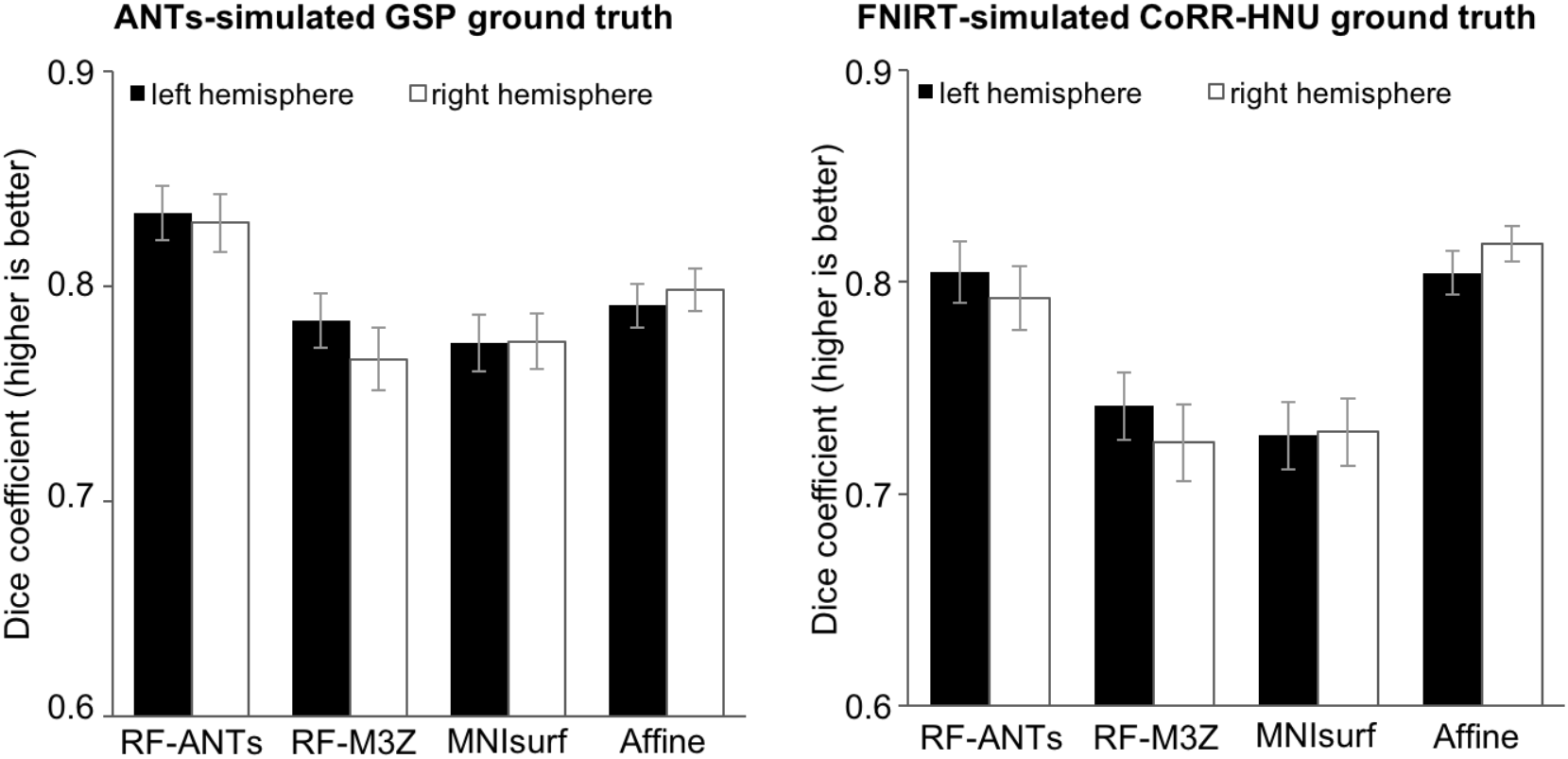
Dice of winner-takes-all fsaverage parcellation projected to MNI152 space, compared against (left) ANTs-simulated GSP “ground truth” and (right) FNIRT-simulated CoRR-HNU “ground truth”. Bars represent Dice coefficient averaged across all 74 segmentation labels within left hemisphere (black) and right hemisphere (white). Error bars correspond to standard errors.

To summarize, RF-ANTs performed the best, although Affine also performed surprisingly well when FNIRT-simulated CoRR-HNU “ground truth” was considered (Figure 13 right). Visual inspection of Figure 12A suggests that the projected cortical ribbon for Affine did not match well to the MNI152 cortical ribbon. However, the dilation within the loose cortical mask appeared to compensate for the poor mapping (Figure 12B), leading to a competitive dice score (Figure 13 right).

### Registration fusion convergence

Figure 14 shows the average NAD metric within each hemisphere for projecting ANTs-derived GSP MNI152 probabilistic maps to fsaverage space, plotted against the number of subjects used to construct the RF mappings. The NAD values start converging after about 100 subjects. When more than 300 subjects are used, the NAD values are mostly stable, although the right hemisphere for RF-ANTs seem to require more subjects to converge. This suggests that the use of 745 training subjects is sufficient to guarantee the quality of the RF mappings. Nevertheless, the RF mappings that we have made publicly available, utilize the entire GSP dataset to construct the mappings.

**Figure 14.**
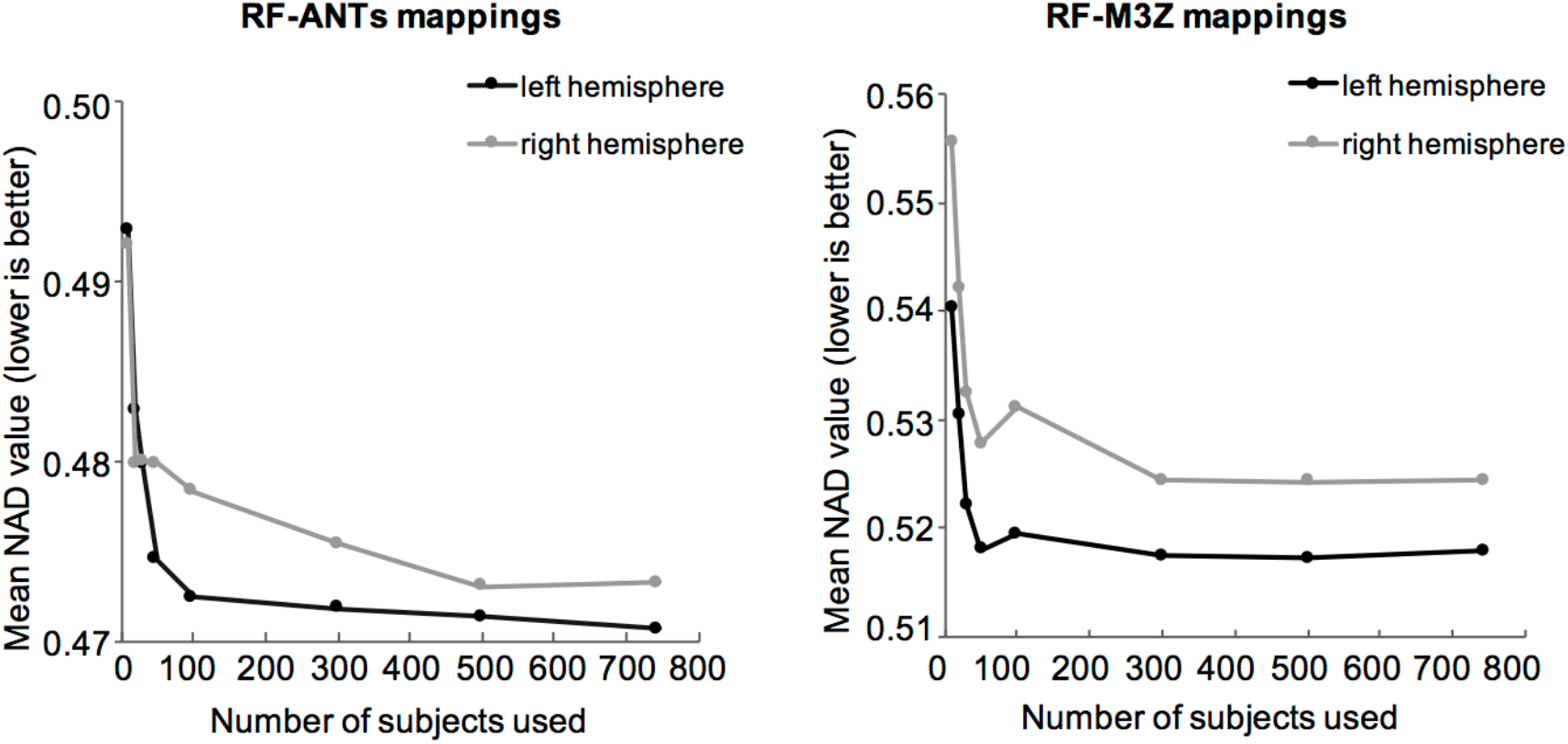
Normalized Absolute Difference (NAD) of ANTs-derived GSP MNI152 probabilistic maps projected to fsaverage space as a function of the number of subjects used to create the RF mappings. (Left) RF-ANTs. (Right) RF-M3Z. NADs were averaged across all 74 probabilistic maps within left hemisphere (black) and right hemisphere (gray), and across all subjects in GSP test set. Results converge after about 300 subjects, although the right hemisphere for RF-ANTs seems to require more subjects to converge.

### Colin27-to-fsaverage and fsaverage-to-Colin27 projections

Figures S1 to S3 show the Colin27-to-fsaverage projection results. The results are largely consistent with the MNI152-to-fsaverage results. In the case of the GSP test set (using ANTs-derived volumetric probabilistic maps), RF-ANTs was the best (p < 0.01 corrected). RF-M3Z and Colin27surf showed comparable performance and were both significantly better than Affine (p < 0.01 corrected). In the case of the CoRR-HNU dataset (using FNIRT-derived Colin27 volumetric probabilistic maps), RF-ANTs was also the best (p < 0.01 corrected). RF-M3Z was the second best (p < 0.01 corrected), followed by Affine (p < 0.01 corrected). Hemispheric differences within each approach were not statistically significant (all p > 0.1). To summarize, RF-ANTs always performed the best.

Figures S4 to S6 show the fsaverage-to-Colin27 projection results. The results are largely consistent with the fsaverage-to-MNI152 results. In the case of the GSP test set (using ANTs-derived “ground truth”), RF-ANTs was the best (p < 0.01 corrected). While Colin27surf showed better performance than RF-M3Z (p < 0.01 corrected), both of them showed (statistically) comparable performance with Affine. On the other hand, in the case of the CoRR-HNU dataset (using FNIRT-derived “ground truth”), both RF-ANTs and RF-M3Z showed (statistically) comparable performance with Affine. Nevertheless, RF-ANTs showed better performance than RF-M3Z (p < 0.01 corrected). Colin27surf performed the worst (p < 0.01 corrected). To summarize, RF-ANTs performed the best, although Affine also performed surprisingly well when FNIRT-simulated CoRR-HNU “ground truth” was considered. Similar to the previous section, the dilation within the loose cortical mask compensated for the actually poor Affine mapping.

## Discussion

In this paper, various approaches (Affine, MNIsurf/Colin27surf, RF-M3Z and RF-ANTs) for mapping between MNI/Colin27 and fsaverage were quantitatively evaluated. RF-ANTs performed the best.

Our results showed that RF-ANTs compared favorably with RF-M3Z, MNIsurf/Colin27surf and Affine even when FSL FNIRT was used to set up the evaluations using a dataset different from the one utilized to derive RF-ANTs. This suggests that if a different software (other than ANTs) was used to register data to MNI152 space, it would still be preferable to use RF-ANTs (rather than RF-M3Z, MNIsurf or Affine) to map the resulting data to fsaverage space. Nevertheless, to achieve best performance, if SPM was used to register data to MNI152 space, then it would probably be the most optimal to generate a new set of RF transformations using SPM.

One potential concern is that the RF mappings are expensive to create because it requires registering a large number of subjects. However, we note that this is a one-time cost. To alleviate this one-time cost, RF-M3Z and RF-ANTs mappings generated using all 1490 GSP subjects are available at https://github.com/ThomasYeoLab/CBIG/tree/master/stable_projects/registration/Wu2017_RegistrationFusion. The code to replicate the mappings or generate new mappings can be found at the same repository.

Recent work has proposed integrating surface-based and volumetric registration to obtain the advantages of each (Joshi et al., 2009; Postelnicu et al., 2009; Zollei et al., 2010). These combined-volume-surface registration algorithms either used both cortical features and volumetric intensity to drive the alignment simultaneously (Joshi et al., 2009) or used geometric information from a surface-based warp to initialize the volumetric alignment (Postelnicu et al., 2009; Zollei et al., 2010). However, these methods were usually designed with inter-subject registration in mind. The combined-volume-surface registration algorithms can be used to create a joint surface-volumetric template, in which the surface and volumetric surface coordinate systems are in alignment. However, creating a new coordinate system would not be helpful for researchers with existing data in MNI152/Colin27 and fsaverage coordinate systems.

The RF approaches might also potentially benefit from improving the registration between subjects and the common coordinate systems (e.g., MNI152). As the RF approaches can be easily adapted to new registration methods, future work can explore more variants of the RF approach. For example, by restricting the registration between subjects’ native space and MNI152/Colin27 to geodesic paths in an anatomical manifold (Hamm et al., 2010), we might be able to generate a better final mapping.

In summary, the RF-ANTs projections between MNI152/Colin27 and fsaverage worked surprisingly well. For example, the projected anatomical structures fitted the ground truth boundaries very well (Figure 4), although there were clear, but minor mis-registrations across sulci. The advantage of registration fusion is consistent with the image segmentation literature, which has demonstrated that using multiple registrations for label fusion can improve image segmentation because the multiple registrations capture greater inter-subject variability and protects against occasional registration failures (Heckemann et al., 2006; Aljabar et al., 2009; Collins and Pruessner, 2010; Sabuncu et al., 2010; Wang et al., 2013; Iglesias and Sabuncu, 2015).

Overall, we believe that the RF approach is useful for projecting between volume and surface coordinate systems. However, we emphasize that the best way of mapping data to fsaverage is by registering subjects directly to fsaverage, while the best way of mapping data to MNI152/Colin27 is by registering subjects directly to the corresponding volumetric template. When data in individuals’ native spaces are available, researchers should not use the RF approaches to project individuals’ data between MNI152/Colin27 and fsaverage for convenience. The RF approaches evaluated in this paper can be considered when the optimal approach is not possible (e.g., when running FreeSurfer on individual subjects is not possible). Furthermore, care must be taken when interpreting results. For example, when describing MNI152 results that have been projected to fsaverage for visualization, it is important to verify that the description is consistent with the original volumetric data in MNI152 space.

## Conclusion

In this paper, we compared various approaches for mapping between MNI152/Colin27 volumetric and fsaverage surface coordinate systems. We found that a new implementation of the RF approach (Buckner et al., 2011; Yeo et al., 2011), RF-ANTs, performed the best. Nevertheless, it is worth noting that the most optimal approach for mapping data to a particular coordinate system (e.g., fsaverage) is to register individual subjects directly to the coordinate system, rather than via another coordinate system. However, in scenarios where the optimal approaches are not possible (e.g., mapping previously published results from MNI152 to fsaverage), we recommend using RF-ANTs. The RF approach can be easily adapted for other volumetric and surface coordinate systems. Code and transformations from this paper can be found at https://github.com/ThomasYeoLab/CBIG/tree/master/stable_projects/registration/Wu2017_RegistrationFusion.

## Acknowledgement

We would like to thank Bin Bin Tang for assistance with testing several algorithms. This work was supported by Singapore MOE Tier 2 (M0E2014-T2-2-016), NUS Strategic Research (DPRT/944/09/14), NUS SOM Aspiration Fund (R185000271720), Singapore NMRC (CBRG/0088/2015), NUS YIA and the Singapore National Research Foundation (NRF) Fellowship (Class of 2017). Our computational work for this article was partially performed on resources of the National Supercomputing Centre, Singapore. Data were also provided by the Brain Genomics Superstruct Project of Harvard University and the Massachusetts General Hospital (Principal Investigators: Randy Buckner, Joshua Roffman, and Jordan Smoller), with support from the Center for Brain Science Neuroinformatics Research Group, the Athinoula A. Martinos Center for Biomedical Imaging, and the Center for Human Genetic Research. Twenty individual investigators at Harvard and MGH generously contributed data to the overall project. Our research also utilized resources provided by the Center for Functional Neuroimaging Technologies, P41EB015896 and instruments supported by 1S10RR023401, 1S10RR019307, and 1S10RR023043 from the Athinoula A. Martinos Center for Biomedical Imaging at the Massachusetts General Hospital. SBE is supported by the National Institute of Mental Health (R01-MH074457), the Helmholtz Portfolio Theme “Supercomputing and Modeling for the Human Brain” and the European Union’s Horizon 2020 Research and Innovation Programme under Grant Agreement No. 7202070 (HBP SGA1). BF is supported by the National Institute for Biomedical Imaging and Bioengineering (P41EB015896, 1R01EB023281, R01EB006758, R21EB018907, R01EB019956), the National Institute on Aging (5R01AG008122, R01AG016495), the National Institute of Diabetes and Digestive and Kidney Diseases (1-R21-DK-108277-01), the National Institute for Neurological Disorders and Stroke (R01NS0525851, R21NS072652, R01NS070963, R01NS083534, 5U01NS086625). Additional support was provided by the NIH Blueprint for Neuroscience Research (5U01-MH093765), as part of the multi-institutional Human Connectome Project. In addition, BF has a financial interest in CorticoMetrics, a company whose medical pursuits focus on brain imaging and measurement technologies. BF’s interests were reviewed and are managed by Massachusetts General Hospital and Partners HealthCare in accordance with their conflict of interest policies.

## Supplementary figures

**Figure S1.**
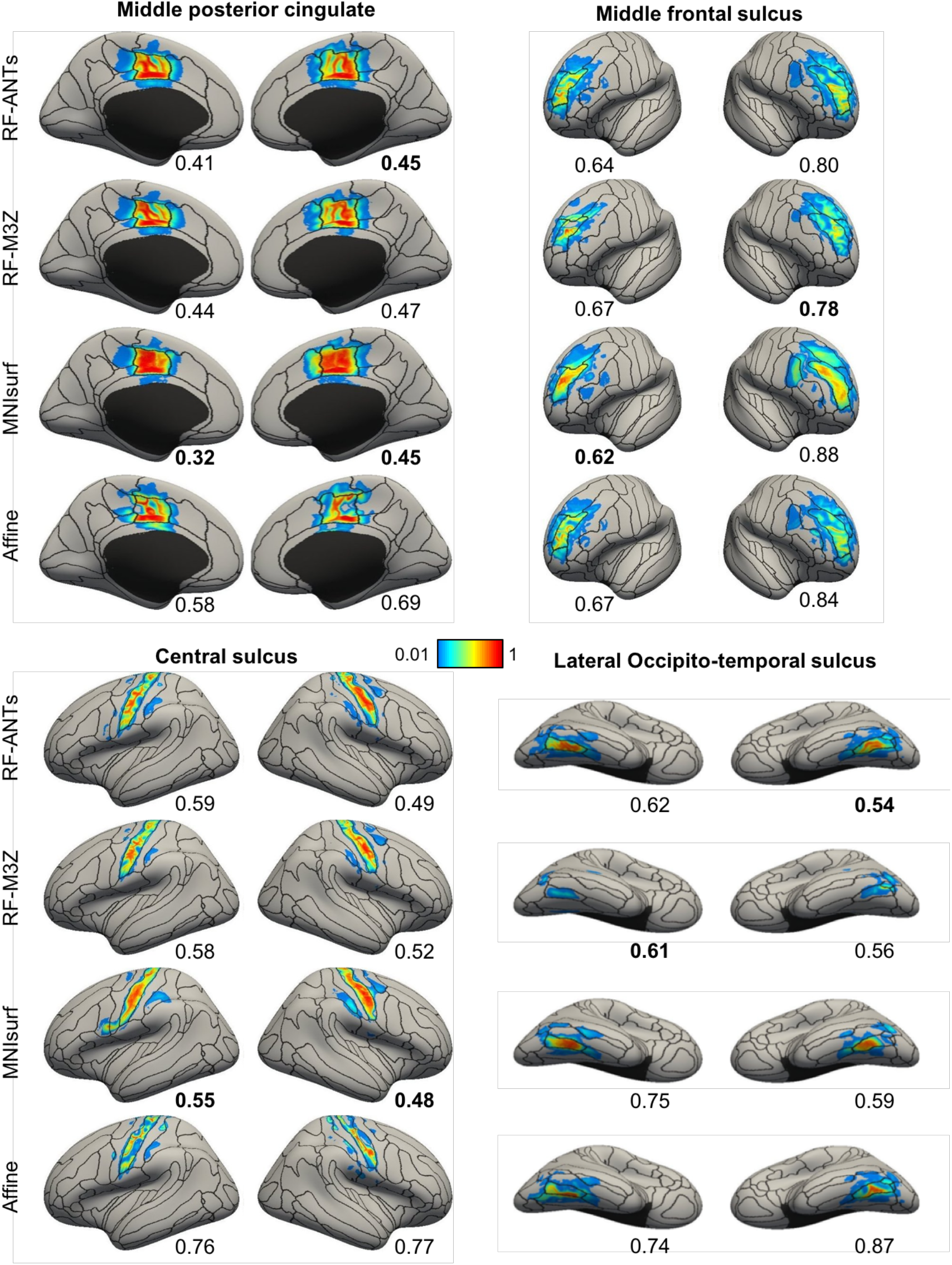
Visualization of ANTs-derived Colin27 probabilistic maps projected to fsaverage surface space in the GSP test set. Four representative structures are shown. Black boundaries correspond to the “ground truth” winner-takes-all parcellation. The value below each cortical surface shows the Normalized Absolute Difference (NAD) between projected probabilistic map and “ground truth” probabilistic map, where a smaller value indicates better performances. Best NAD for each region is **bolded**.

**Figure S2.**
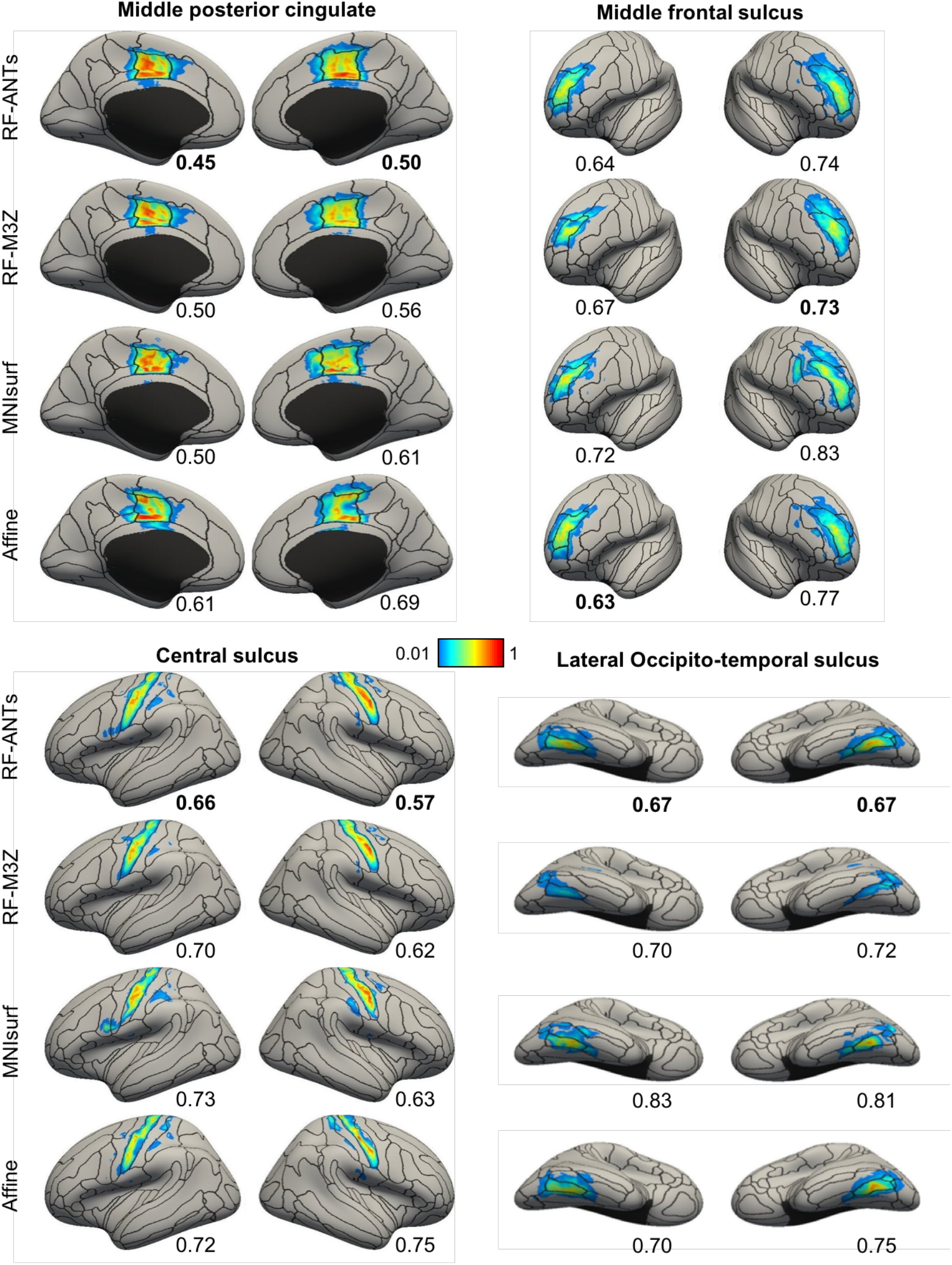
Visualization of FNIRT-derived Colin27 probabilistic maps projected to fsaverage surface space in the CoRR-HNU dataset. Four representative structures are shown. Black boundaries correspond to the “ground truth” winner-takes-all parcellation. The value below each cortical surface shows the Normalized Absolute Difference (NAD) between projected probabilistic map and “ground truth” probabilistic map, where a smaller value indicates better performances. Best NAD for each region is **bolded**.

**Figure S3.**
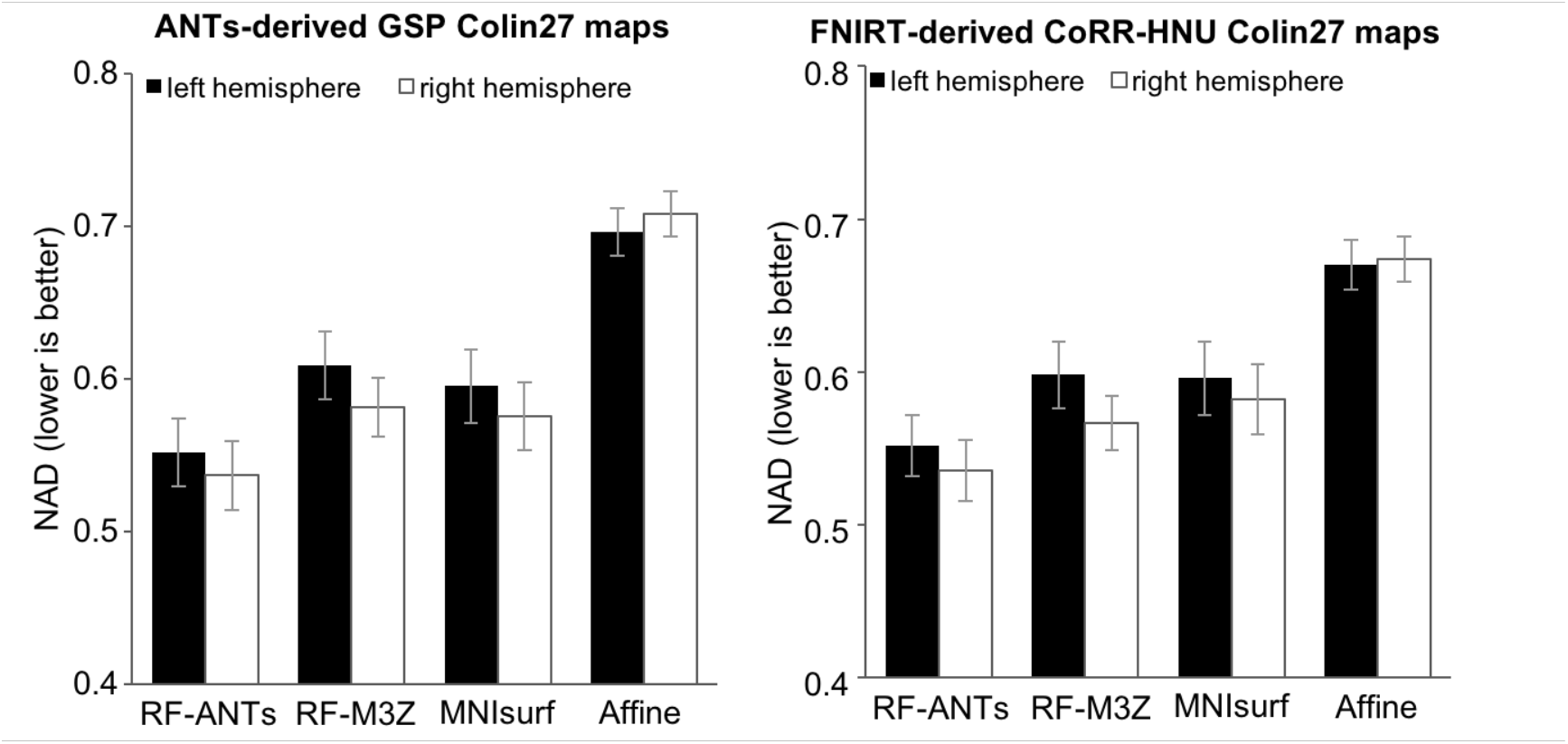
Normalized Absolute Difference (NAD) of Colin27 probabilistic maps projected to fsaverage surface space. (Left) Results for ANTs-derived GSP MNI152 probabilistic maps. (Right) Results for FNIRT-derived CoRR-HNU Colin27 probabilistic maps. The bars represent the NADs averaged across all 74 probabilistic maps within left hemisphere (black) and right hemisphere (white). Error bars correspond to standard errors across the 74 anatomical structures. Overall, RF-ANTs performed the best. Numerically, it may seem that the right hemisphere show smaller NAD values for the nonlinear methods and larger NAD value for Affine, compared to the left hemisphere. However, the difference were not significant (all p > 0.1).

**Figure S4.**
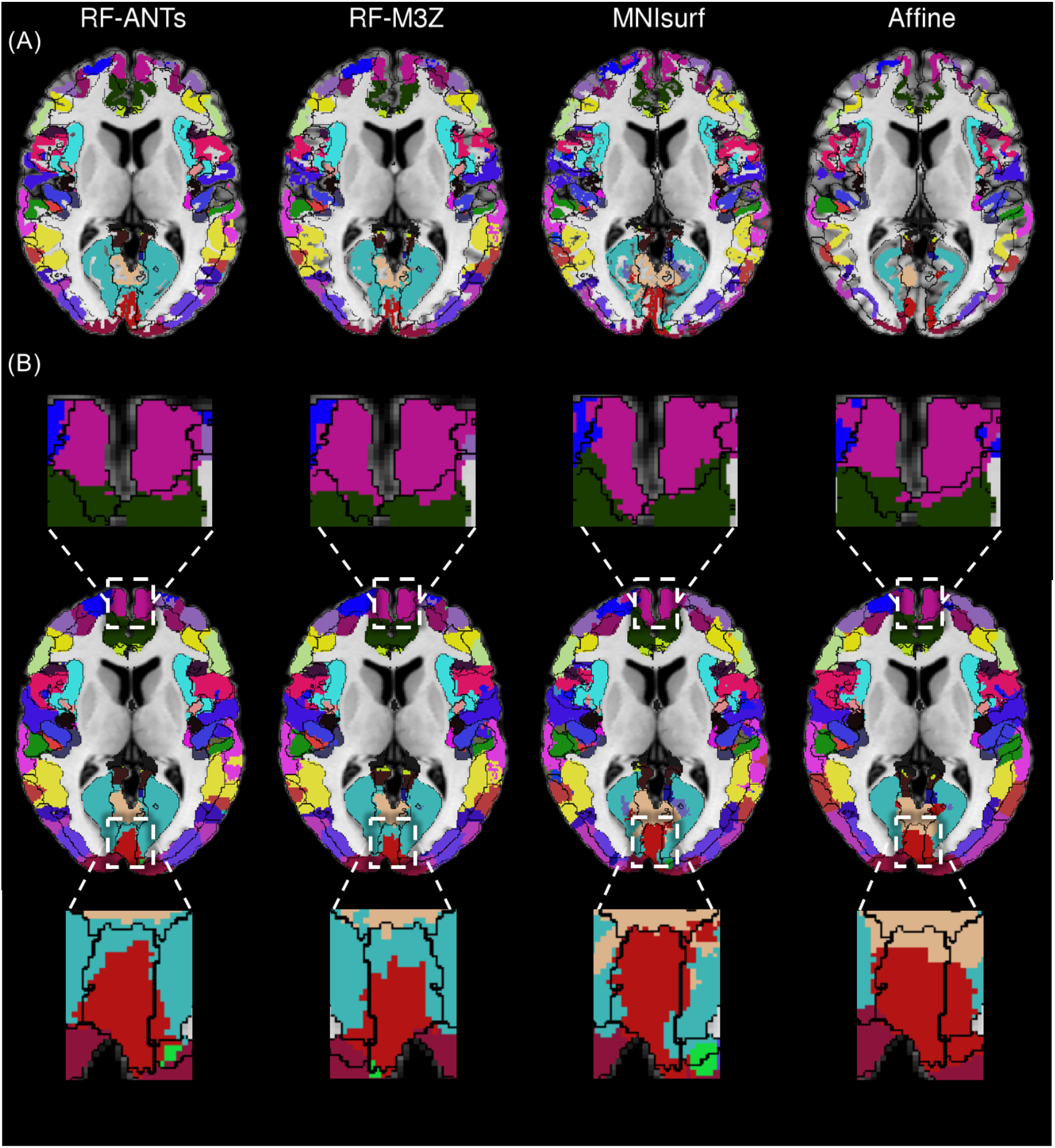
Winner-takes-all fsaverage parcellation projected to Colin27 volumetric space with ANTs-simulated “ground truth” (black boundaries) in the GSP test set. (A) Projections before dilation within loose cortical mask. (B) Projections after dilation within loose cortical mask.

**Figure S5.**
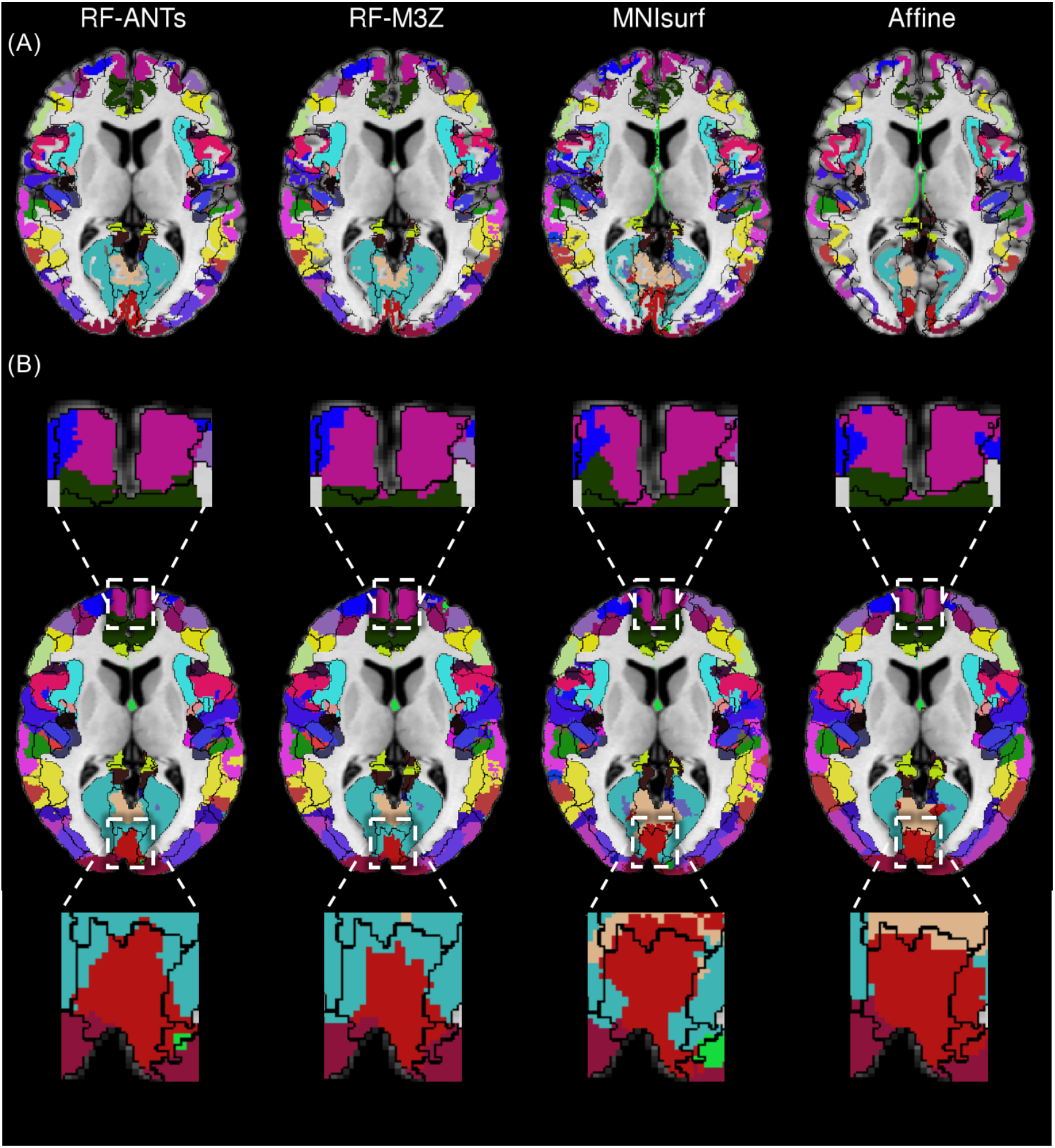
Winner-takes-all fsaverage parcellation projected to Colin27 volumetric space with FNIRT-simulated “ground truth” (black boundaries) in the CoRR-HNU dataset. (A) Projections before dilation within loose cortical mask. (B) Projections after dilation within loose cortical mask.

**Figure S6.**
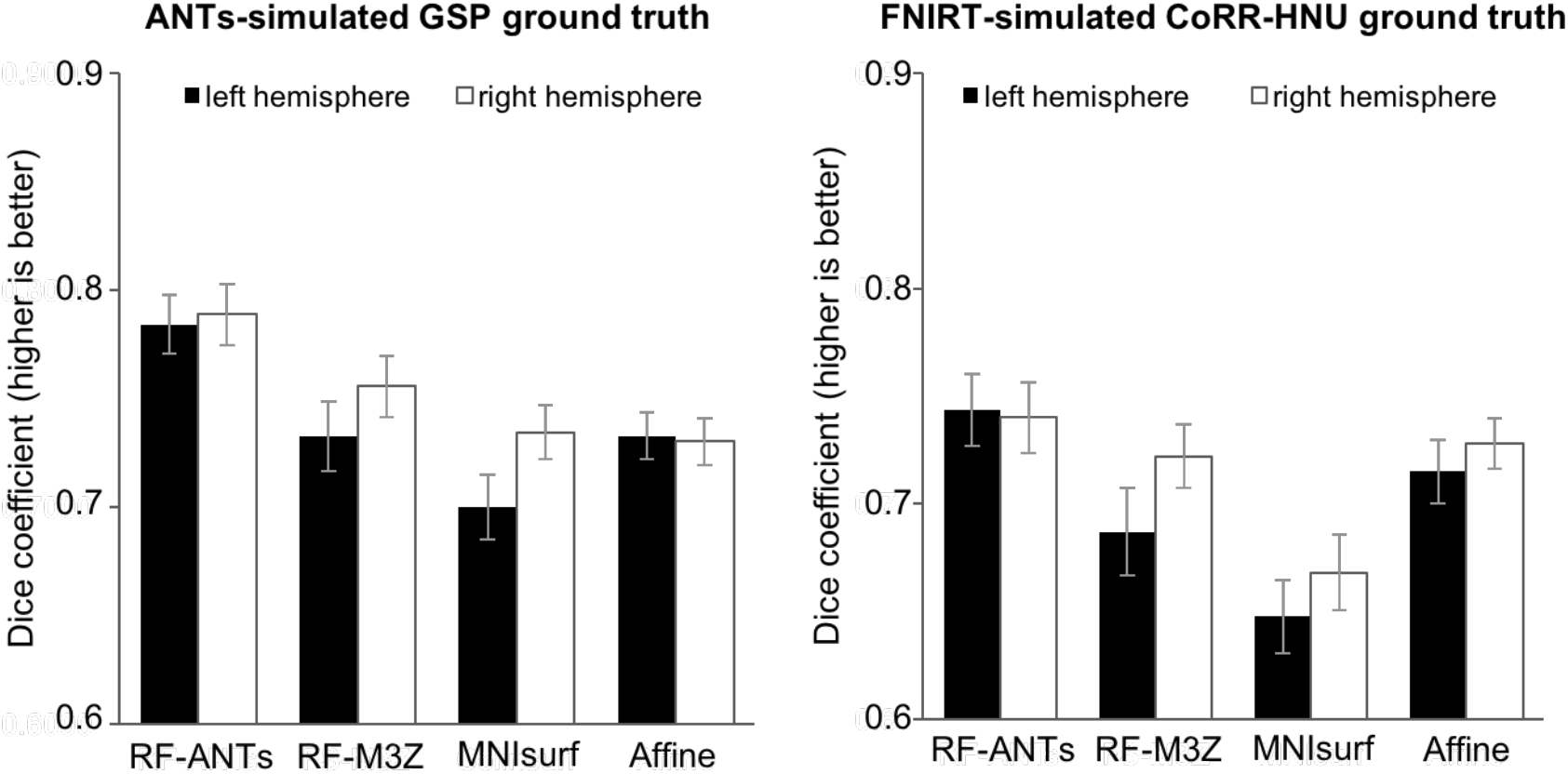
Dice of winner-takes-all fsaverage parcellation projected to Colin27 space, compared against (left) ANTs-simulated GSP “ground truth” and (right) FNIRT-simulated CoRR-HNU “ground truth”. Bars represent Dice coefficient averaged across all 74 segmentation labels with left hemisphere (black) and right hemisphere (white). Error bars correspond to standard errors.

## Footnotes

1 Note that this internal volumetric space is different from fsaverage volume.

